# Systematic elucidation and pharmacologic targeting of non-oncogene dependencies in imatinib-resistant gastrointestinal stromal tumor

**DOI:** 10.1101/2025.10.12.681609

**Authors:** Prabhjot S. Mundi, Adina Grunn, Arsenije Kojadinovic, Sergey Pampou, Charles Karan, Ronald Realubit, Cristina I. Caescu, Hanina Hibshoosh, Mahalaxmi Aburi, Mariano J. Alvarez, Matthew Ingham, Denisse Evans, Sara Rothschild, Gary K. Schwartz, Andrea Califano

## Abstract

Treatment of gastrointestinal stromal tumor (GIST) with imatinib and other KIT-targeting drugs has improved outcomes significantly. However, most patients with advanced GIST eventually develop imatinib resistance and succumb to disease. We have developed mutation-agnostic, network-based methodologies to systematically elucidate and pharmacologically target Master Regulator (MR) proteins—critical non-oncogene dependencies—in cancer cells. Unsupervised, MR-based clustering of 34 GIST patient tumor samples produced two clusters, one of which contained all imatinib-resistant tumors. Analysis of 9 single-cell RNA profiles of high-risk GIST revealed that tumors with clinical progression on imatinib harbored large subpopulations enriched for the MR-activity signature of imatinib-resistant tumors, while tumors with resistance-associated mutations but without overt progression showed smaller, variably sized enriched subpopulations. High-throughput profiling of transcriptional responses by two GIST cell lines to FDA-approved and late-stage experimental drugs identified six candidate drugs that reversed the MR activity of imatinib-resistant GIST. Predictions were validated in two imatinib-resistant, patient-derived xenograft (PDX) models. The top prediction, linifanib, induced marked tumor growth inhibition in both PDXs across a wide dose range; selinexor and selumetinib were also effective compared to imatinib. We confirmed *in vivo* MR-activity reversal by these drugs, but not by ineffective drugs.

## Introduction

The oncogene addiction paradigm has been central to personalized oncology for the past three decades^1^, with gastrointestinal stromal tumor (GIST) representing one of its earliest successful clinical applications. In this archetype, tumors are highly dependent on the aberrant activity of a singular protein encoded by a recurrently mutated oncogene.

GIST is a malignancy of mesenchymal origin arising within the gastrointestinal tract, most frequently the stomach or small intestine, and is one of the most common subtypes of visceral sarcoma^2^. The putative lineage of origin of GIST is the interstitial cells of Cajal, pacemaker cells within the inner circular layer of the muscularis mucosa that propagate intestinal motility, based on the observation of near universal membranous expression of DOG-1 and c-KIT in these tumors^3–6^. While some localized GISTs can be cured with surgery alone, higher risk GISTs, characterized by larger tumor size, high mitotic rate, non-stomach location, and intraperitoneal tumor rupture, frequently recur and metastasize, becoming incurable. In the metastatic setting, GISTs are resistant to cytotoxic chemotherapy and radiation^7^.

Approximately 70-80% of GISTs harbor activating mutations in the receptor tyrosine kinase (RTK) *KIT*, most commonly deletions in exon 11 encoding the juxtamembrane domain, with mutually exclusive mutations in the related RTK *PDGFRA* occurring in an additional 5-10%^3,8^. Somatic or germline mutations in *BRAF*, *NF1*, or succinate dehydrogenase complex subunit genes occur in a sizeable proportion of the remaining *KIT/PDGFRA*-wildtype GISTs^9–11^.

Imatinib, a small molecule tyrosine kinase inhibitor (TKI) with selectivity for c-KIT and PDGFR-alpha, reduces relapse risk in the adjuvant setting and is first line therapy for patients with metastatic GIST. Imatinib has demonstrated objective response rates around 54% by RECIST criteria^12^ and extended median overall survival for metastatic disease from 18 to 56 months^13,14^. The response to imatinib, however, is highly heterogenous; 15% of GISTs present with primary resistance, mainly in *KIT*-wildtype or *KIT* exon 9 mutated tumors, while a similar percentage of patients have durable disease control for over five years on imatinib^15^. Most patients fall somewhere in between, with median progression free survival (mPFS) of 18 to 24 months.

Unfortunately, acquired resistance to imatinib eventually occurs in the majority of patients with advanced GIST, and is in part attributed to secondary mutations in *KIT*, including amplification, loss of heterozygosity, and point mutations in the activation loop (exons 17/18) or the ATP-binding pocket (exons 13/14) that reduce imatinib binding^16,17^. Other potential mechanisms of resistance include activation of parallel or downstream kinase pathways^18–20^, pharmacokinetic changes resulting in decreased drug levels over time mediated by upregulation of ABC transporters in the intestine and increased erythrocyte uptake^21–23^, as well as potential epigenetic cell adaptation. Two multi-kinase inhibitors, sunitinib and regorafenib, are approved in the second- and third-line setting, although the duration of clinical benefit from these agents is progressively shorter, with mPFS around 6.3 and 4.8 months, respectively^24,25^. More recently, the next-generation highly selective c-KIT and PDGFR inhibitors ripretinib and avapritinib were also approved based on their activity against mutations that confer resistance to imatinib including *KIT* exon 17 and *PDGFRA* D842V, although objective response rates are low^26,27^.

***OncoTarget*** and ***OncoTreat*** are transcriptome-based, mutation-agnostic methodologies that predict drugs targeting Master Regulator proteins (MRs)—non-oncogene dependencies and mechanistic determinants of tumor cell state. These have been validated both preclinically^28,29^ and clinically^30,31^. Both leverage the VIPER algorithm^32^, which identifies aberrantly activated and inactivated proteins, based on the differential expression of their transcriptional targets. The most aberrantly activated and inactivated proteins represent candidate MRs capable of eliciting tumor-specific essentiality^33,34^ or synthetic lethality^35,36^. Candidate MRs, collectively, form tightly autoregulated modules that implement highly specific on/off switches controlling cancer cell transcriptional state^37^. VIPER has identified mechanisms of tumorigenesis, progression, and drug resistance in glioma^38^, leukemia^39^, lymphoma^40^, prostate^35^ and breast cancer^41^, among others. Once candidate MR dependencies are identified, *OncoTarget* prioritizes clinically relevant drugs representing high-affinity MR inhibitors, while *OncoTreat* predicts drugs capable of reversing the activity of the entire repertoire of aberrant MR activity^28,33^. The latter is accomplished by analyzing post-drug perturbation profiles—representing the transcriptional response of high-fidelity cell lines—to assess differential MR protein activity in drug vs. vehicle control-treated cells^42,43^.

Herein, we describe the application of these methodologies to identify novel MRs and MR-targeting drugs for the treatment of imatinib-resistant GIST. For this purpose, we collaborated with the Life Raft Group, a patient advocacy organization that has compiled a large GIST tumor bank, with extensive annotation of clinicopathological features and clinical outcomes. We performed transcriptomic profiling of GIST tumors at various stages of disease trajectory, including: initial diagnosis, on-treatment with imatinib in the adjuvant or metastatic setting, and after the development of clinical progression on imatinib and in some cases after resistance to multiple subsequent treatments such as sunitinib and regorafenib.

Tumor-specific MRs—identified by unbiased VIPER inference—emerged as highly consistent across imatinib-resistant tumors, representing candidate mechanistic determinants of imatinib-resistant vs. sensitive state. Analysis of a separate published cohort^44^ of single-cell RNASeq profiles of high-risk GIST identified large tumor subpopulations enriched for this same MR-activity signature of resistance in tumors that had demonstrated clinical progression on imatinib, and smaller, variable-sized subpopulations in tumors harboring mutations associated with imatinib resistance but prior to the development of frank clinical progression. We identified six candidate drugs for follow up *in vivo* validation, based on their highly significant predicted MR targeting of the imatinib-resistant state, two by *OncoTarget* and four by *OncoTreat* analysis. The latter were based on pre- and post-drug perturbation RNA profiles generated in two established GIST cell lines presenting differential imatinib sensitivity. We then collaborated with Crown Bioscience, to identify—among 18 available GIST PDX models—the two that most significantly recapitulated the MR activity signature of imatinib-resistant patient tumors, as optimal models for drug efficacy evaluation. Linifanib, the top predicted drug, induced tumor growth inhibition in both PDXs, across a wide dose range, including tumor regression at the highest dose level. Other predicted drugs also compared favorably to imatinib. We further demonstrate proof-of-concept through on-treatment biopsy profiling, and demonstrate *in vivo* reversal of MR activity with effective drugs, but not with ineffective drugs.

We discuss the implications of our findings for imatinib-resistant GIST, including clinical trial validation, single-cell RNASeq profiling to characterize intra-tumoral heterogeneity and resistance at the subpopulation level, and the potential use of an *OncoTreat-*based framework to dynamically target evolving drug resistance, applicable across cancer types.

## Results

### Characterizing Master Regulators of Imatinib resistance

We collaborated with the Life Raft Group (LRG) patient advocacy organization, to profile GIST patient tumor samples obtained at various stages of disease, spanning diagnosis, on imatinib treatment, imatinib-resistant disease, and multi-drug progression. The LRG maintains a GIST patient registry and a companion tissue bank, which links clinical information in the registry to biological samples.

RNASeq profiles were generated from 34 GIST samples from the registry that met quality control standards (**Table S1**), including 10 samples obtained after progression to imatinib-resistant disease. **Table S2** provides associated demographic and clinicopathologic annotation for all samples, including *KIT/PDGFRA* genotype, relative timing of sample acquisition, time on various treatments, and progression and survival outcomes when available. Differential protein activity was assessed by metaVIPER^45,46^, using the centroid of the dataset as reference for differential expression analysis, see **Methods**.

Unsupervised, protein activity-based consensus clustering (see **Methods**^47,48^) identified k=2 as the optimal cluster solution (**Figure 1A**), producing the lowest connectivity and highest silhouette index (**Figure 1B-C**). Of the 16 samples in the first cluster (***C*_1_**), 10 represented imatinib-resistant cases while all samples in the second cluster (***C*_2_**) were imatinib sensitive or naïve to treatment (Fisher’s exact two-tailed *p* = 0.0001) (**Figure 1A**). Of the 6 samples in ***C*_1_**that are not imatinib-resistant, 4 were not metastatic at diagnosis and had not developed recurrent disease, and potentially represent aggressive cases that were fortuitously cured with early surgical intervention. While protein activity and cluster analyses were performed agnostic to genotype and clinicopathological features, ***C*_1_** also contained all n=3 tumors harboring *PDGFRA* exon 18 (D842V) mutations and all n=3 *KIT/PDGFRA* wildtype tumors, which are known to confer primary resistance to imatinib^49^. Based on this cluster solution (**Figure 1A**), only four of the 34 samples presented with ambiguous cluster placement [i.e., not placed in the same cluster group across 10,000 iterations run], with three of the four harboring *KIT* exon 9 alterations, which are associated with decreased efficacy of imatinib.

**Figure 1.**
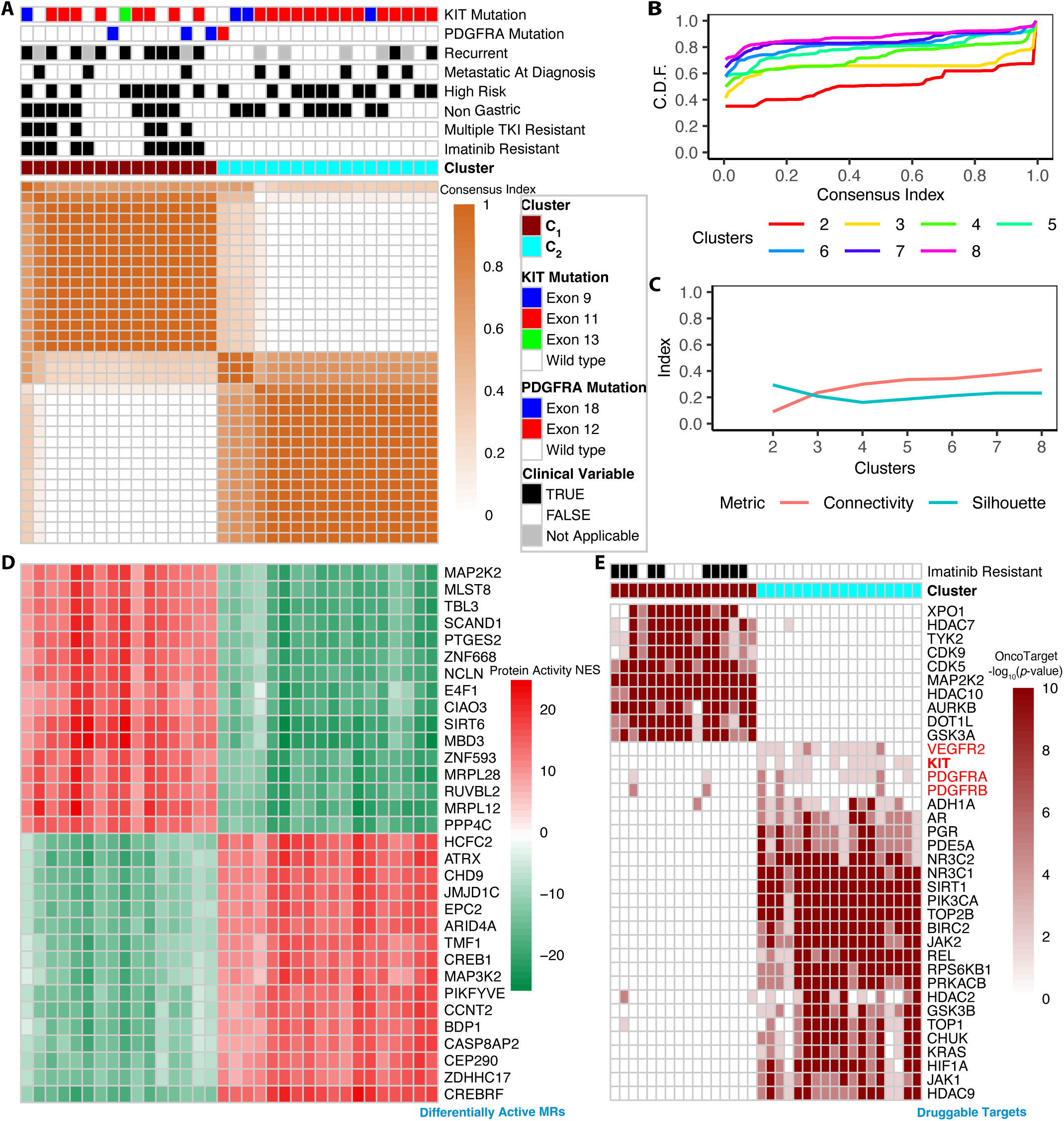
Protein activity profiling of Life Raft GIST registry patient tumors. Bulk RNASeq profiling of 34 unique patient samples from the Life Raft registry was completed, including 10 from patients whose cancer had relapsed or progressed while receiving imatinib therapy. MetaVIPER analysis was used to infer differential protein activity in each tumor. ***(A)*** Unsupervised consensus clustering of tumors was performed on the protein activity signatures, using a partition around medoids (PAM) approach, with n=10,000 iterations. The optimal solution to consensus clustering occurred with k=2 clusters. The symmetric consensus matrix heatmap for k=2 demonstrates that most sample pairings either always cluster together (consensus index ∼1, dark brown) or never cluster together (consensus index ∼0, white), with four samples (first column, and three columns in the middle) demonstrating more ambiguous clustering across iterations. Cluster assignment and clinical annotation are provided at the top of the heatmap. ***(B)*** Consensus index distribution for different partition numbers (k=2 to 8). The cumulative distribution for k=2 and k=3 contain the fewest ambiguous sample pairings whose consensus index is between 0.2 and 0.8. ***(C)*** Additional metrics for optimal partitioning using a PAM approach. The connectivity index (the inverse of the average number of in-cluster samples that are a shorter distance than the closest out-of-cluster sample) and silhouette index (the relative difference in average distance between in-cluster samples and the nearest out-of-cluster sample) are optimized (lowest connectivity and highest silhouette) for k=2 clusters. ***(D)*** Heatmap of proteins demonstrating the highest variance in activity between clusters. Analysis considers all proteins (n=7,070) whose activity is inferred by metaVIPER. Columns are organized in the same order as *(**A**)*. ***(E)*** Heatmap of *OncoTarget* analysis, representing the most differentially activated of a panel of 180 directly druggable proteins. Columns are organized in the same order as *(**A**)* and color intensity corresponds to the −log10 (BH-corrected *p*) for protein activation.

Based on these analyses, we proceeded to rank all differentially active proteins by their integrated *p*-value across all ***C*_1_**and ***C*_2_** samples, respectively, using Stouffer’s method. From the n=7,070 proteins whose activity was inferred by metaVIPER, the 25 most activated and 25 most inactivated proteins in ***C*_1_** vs. ***C*_2_**samples (and vice-versa) are shown in **Figure 1D**. Consistent with the sharp clustering solution, the most activated and inactivated proteins were highly consistent across ***C*_1_** and ***C*_2_** samples—including MAP2K2 (MEK2), TBL3, ZNF668, and E4F1 (a key post-translational modulator of p53 activity^50^) as aberrantly activated MRs, and C5orf41 (CREBRF) as an inactivated MR in the majority of ***C*_1_**tumors. This suggests conservation of the mechanisms that preside over imatinib sensitivity and resistance across tumors, with imatinib resistance emerging as a transcriptionally distinct cell state. For simplicity, hereafter we use cluster ***C*_1_** and “imatinib-resistant cluster” interchangeably, with the understanding that additional subtle molecular alterations may exist between the 10 tumors that have developed frank clinical progression on imatinib and the remaining 6 tumors.

Pathway enrichment analysis on the integrated protein activity signature of ***C*_1_** vs. ***C*_2_** cluster, using the MSigDB^51,52^ gene ontology, cancer hallmarks^53^, immunological signatures, and oncogenic signatures is presented in **Figure S1**. In C**_1_**tumors, there is downregulation in MYC-proliferative signaling, metabolic alterations (e.g., downregulation of oxidative phosphorylation pathway), and complex changes in DNA repair, including an overall deactivation in DNA repair pathways (e.g., base excision, nucleotide excision, and mismatch repair) and the p53 pathway, as reflected in the ‘Cancer Hallmarks’ pathways. There are also global alterations in transcriptional machinery, as reflected in strong enrichment in ‘Gene Ontology’ pathways related to DNA binding by transcription factors in ***C*_1_**tumors, and complex alterations in immunological signaling^53,54^ based on analysis of the ‘Immunologic Signatures’ gene sets.

### OncoTarget Drug Prediction for Imatinib-resistant Tumors

The *OncoTarget* algorithm has been developed to identify those MR proteins that are also high-affinity targets of clinically relevant drugs^28,45^. *OncoTarget* (see **Methods**) matches differentially activated MRs (*p* < 10^−5^, Benjamini-Hochberg (BH)-corrected) to a curated target-to-drug list, consisting of 180 high-affinity targets of FDA approved and late-stage experimental (phase 2 and 3) antineoplastic drugs (**Table S3**). The majority of such targets are rarely mutated in cancer, and thus, represent candidate non-oncogene dependencies.

Figure 1E summarizes the metaVIPER-assessed activity of the most statistically significant druggable MRs across the 34 GIST samples, ordered by their integrated *p*-value across each cluster, as well as the activity of c-KIT. The analysis identified MAP2K2 (MEK2), HDAC10, exportin-1 (XPO1), CDK9, and DOT1L as recurrently activated targets (BH-corrected *p* ≤ 10^−5^) across a majority of the 10 imatinib-resistant tumors. Interestingly, c-KIT activity was modestly, yet statistically significantly higher in ***C*_2_** (imatinib-sensitive) vs. ***C*_1_** samples (two sample t-test of protein activity scores, *p* = 2.2 x 10^−16^). Only a few recurrently activated, druggable targets were identified by *OncoTarget*, some of which match to drugs that are difficult to attain for therapeutic study (e.g., pinometostat for DOT1L) or that are considered to be significantly toxic for clinical translation (e.g., the pan-CDK inhibitor dinaciclib for CDK5/9).

### Detecting the emergence of the *C_1_* signature in single cell RNA profiles of high-risk GIST

We analyzed a previously published single-cell (sc) RNASeq dataset of n=9 GIST patient tumors^44^. These include n=2 tumors without demonstrated clinical progression on imatinib and with canonical *KIT* exon 11 [sensitive] mutations only, n=4 tumors that had not yet demonstrated clinical progression but with mutations strongly associated with primary (*PDGFRA* D842V) or acquired imatinib resistance (secondary mutations in *KIT* exons 17/18 that reduce drug binding), and n=3 tumors that had already developed clinical progression on imatinib (see **Table S4** for additional sample annotation). See **Methods** for quality control and malignant cell compartment isolation criteria.

Figure 2, ***top*,** demonstrates a UMAP projection on metaVIPER inferred protein activity of putative malignant cells from each sample. The projection is annotated based on enrichment of the imatinib-resistant cluster ***C_1_*** signature in the metaVIPER protein activity profile of each tumor cell. Larger red dots represent cells strongly enriched in this signature (ImResS+), while larger blue dots represent cells negatively enriched in this signature (ImResS-). The 2 tumors without clinical progression on imatinib and without any treatment-emergent mutations associated with resistance (panels **A,B**) have none or miniscule ImResS+ subpopulations, respectively. The 4 tumors without established clinical progression but with mutations associated with the development of resistance (panels **C,D,E,F**) are observed to have variable-sized ImResS+ subpopulations, ranging from non-existent (**D**) to small (**E**) to substantial (**C,F**). Finally, all 3 tumors with confirmed clinical progression on imatinib (panels **G,H,I**) are observed to have substantial, discrete, ImResS+ subpopulations representing 40.5% to 62.2% of all putative malignant cells.

Interestingly, ImResS+ cells do not demonstrate elevated c-KIT activity relative to their ImResS-counterparts, and in fact are more likely to demonstrate modestly but consistently lower c-KIT activity (displayed underneath the density plots in **Figure 2**, ***bottom***, and also annotated in **Figure S2**, ***top,*** UMAP projections), with Spearman-rank correlation ρ = −0.89. Enrichment for the ImResS signature also does not correlate to *KIT* mRNA differential expression (**Figure S2**, ***bottom***), with Spearman ρ = 0.09 across all samples. In tumors with significant ImResS+ subpopulations, these subpopulations do not appear to correspond to additional inferred copy number alterations (**Figure S3**), including the previously described chromosome 16q deletions that are frequent secondary events in GIST progression^55^ and are inferred for the samples in panels **A,D,E**, with the caveat of the significant limitations of copy number inference from scRNASeq data^56^. The findings potentially support a treatment-emergent imatinib-resistant subpopulation with complex molecular reprogramming that significantly expands by the time clinical resistance develops.

**Figure 2.**
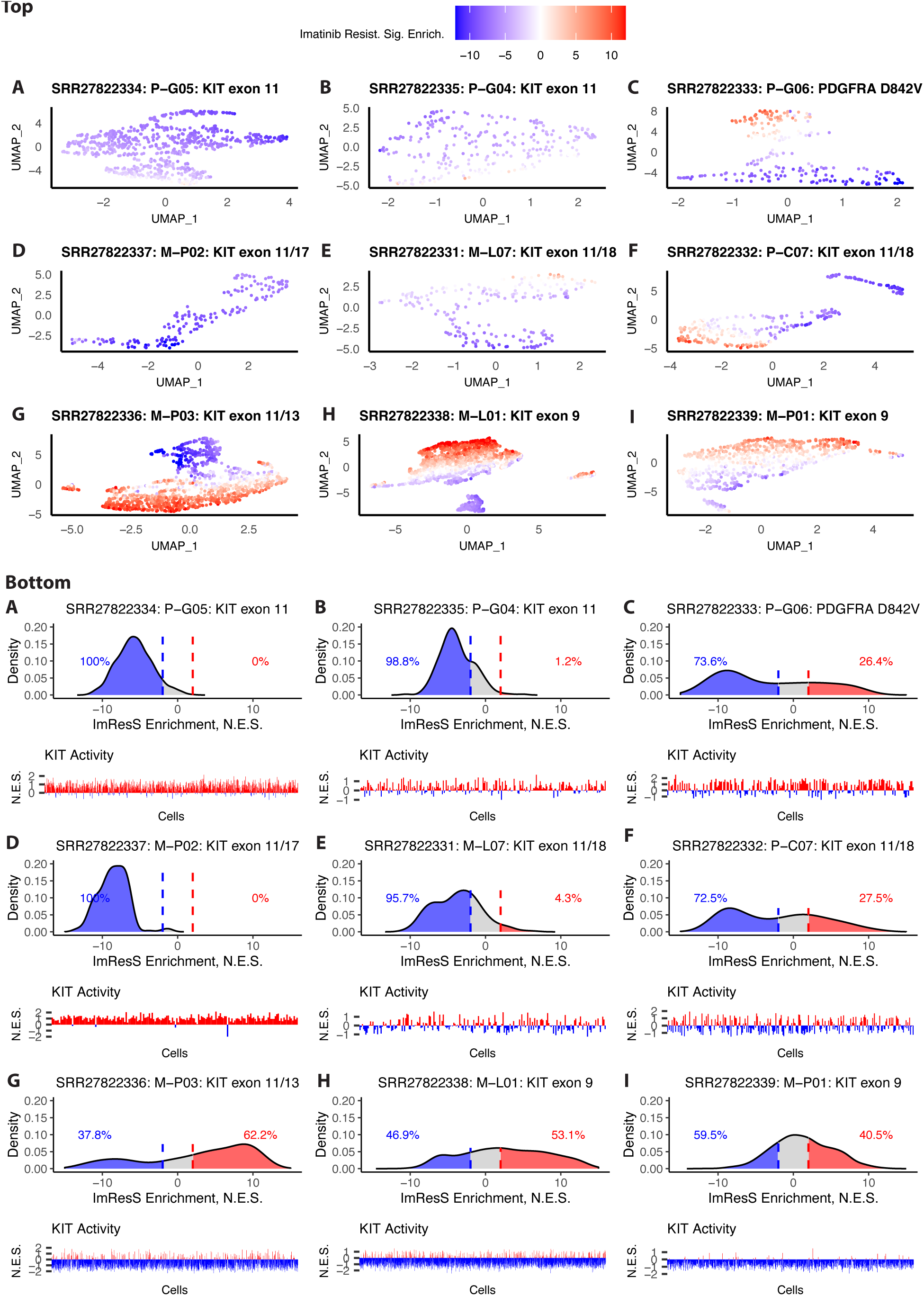
Malignant cell subpopulations enriched for the imatinib-resistant cluster signature in an independent dataset of 9 scRNA profiles of high-risk GIST. (**Top A-I**) A UMAP projection of metaVIPER inferred protein activity signatures (n=7,070 proteins) of single cells within the malignant compartment of each sample, with individual cells further annotated by their enrichment for the imatinib-resistant cluster (**C_1_**) signature, hereafter *ImResS*, identified from analysis of bulk RNA profiles (**Figure 1**). Larger red dots represent cells strongly enriched in this signature (ImResS+), while larger blue dots represent cells negatively enriched in this signature (ImResS-). The two tumors without clinical progression on imatinib and without any treatment-emergent mutations associated with resistance are in panels **A,B**. The four tumors without established clinical progression but with mutations associated with the development of resistance are in panels **C,D,E,F** and are observed to have variable-sized ImResS+ subpopulations. Finally, all three tumors with confirmed clinical progression on imatinib are in panels **G,H,I**. (**Bottom A-I**) Density plots for ImResS enrichment for single cells in each sample are shown in the top portions of each panel, with the blue and red tails highlighting ImResS-cells (normalized enrichment score, NES < −2.0) and ImResS+ cells (NES > 2.0), respectively, further annotated by the percentage of cells in each. The bottom portion of these panels display the c-KIT activity of the corresponding cells ordered left-to-right from lowest to highest ImResS. While the overall differential activity of c-KIT is modest (most NES scores between −1.5 and 1.5), there is consistent anticorrelation between ImResS enrichment and c-KIT activity.

### Generation of Drug Perturbation Profiles and OncoTreat Drug Predictions

*OncoTreat* is an RNA-based assay to predict tumor sensitivity based on a drug’s ability to reverse the overall activity of MR proteins^28,33^ (see **Methods**). It relies on pre- and post-drug perturbation RNA profiles generated in carefully selected *in vitro* models that reasonably recapitulate the patient-specific MR signature(s) of interest, based on the OncoMatch algorithm^28,29^, and VIPER assessment of MR activity change induced by each drug. Drugs are tested at sub-lethal concentrations to reduce confounding from engagement of cell death and stress pathways^28,57,58^. *OncoTreat* uses a conservative statistical significance threshold (*p* ≤ 10^−5^, BH-corrected, by 1-tailed analytic-rank based enrichment analysis — aREA^32^) to identify drugs eliciting activity inversion of the 25 most differentially activated (25↑) and 25 most inactivated (25↓) proteins from each tumor sample, see **Methods**. Notably, *OncoTreat* completely ignores drug sensitivity *in vitro* (in fact, all drugs are profiled at their EC_20_ concentration), and rather uses *in vitro* models only as a means for *de novo* assessment of context-specific drug mechanism (i.e., induced protein activity change).

Among the limited number of available GIST cell line models, we selected GIST-T1 and GIST430^19,59^ to generate pre- and post-drug perturbation RNA profiles. Both GIST-T1 and GIST430 were derived from patient tumors with canonical *KIT* exon 11 deletions—associated with initial imatinib sensitivity, but GIST430 has a secondary, cis-allelic, *KIT* exon 13 mutation^19,59^. GIST-T1 is highly imatinib-sensitive with IC_50_ < 0.1*u*M and GIST430 has been reported as relatively imatinib-resistant, although it must be cultured with low dose imatinib (0.1*u*M) to maintain a resistant state. OncoMatch analysis^28,29^ (see **Methods**) confirmed that both cell lines recapitulate the MR signature of ***C*_1_**and ***C*_2_** samples at our statistical significance threshold (*p* ≤ 10^−5^, BH-corrected) (**Figure 3A**). GIST430 provides a modestly stronger match for ***C*_1_**samples (*p* = 4.81e−07) compared to GIST-T1 (*p* = 3.14e−06).

**Figure 3.**
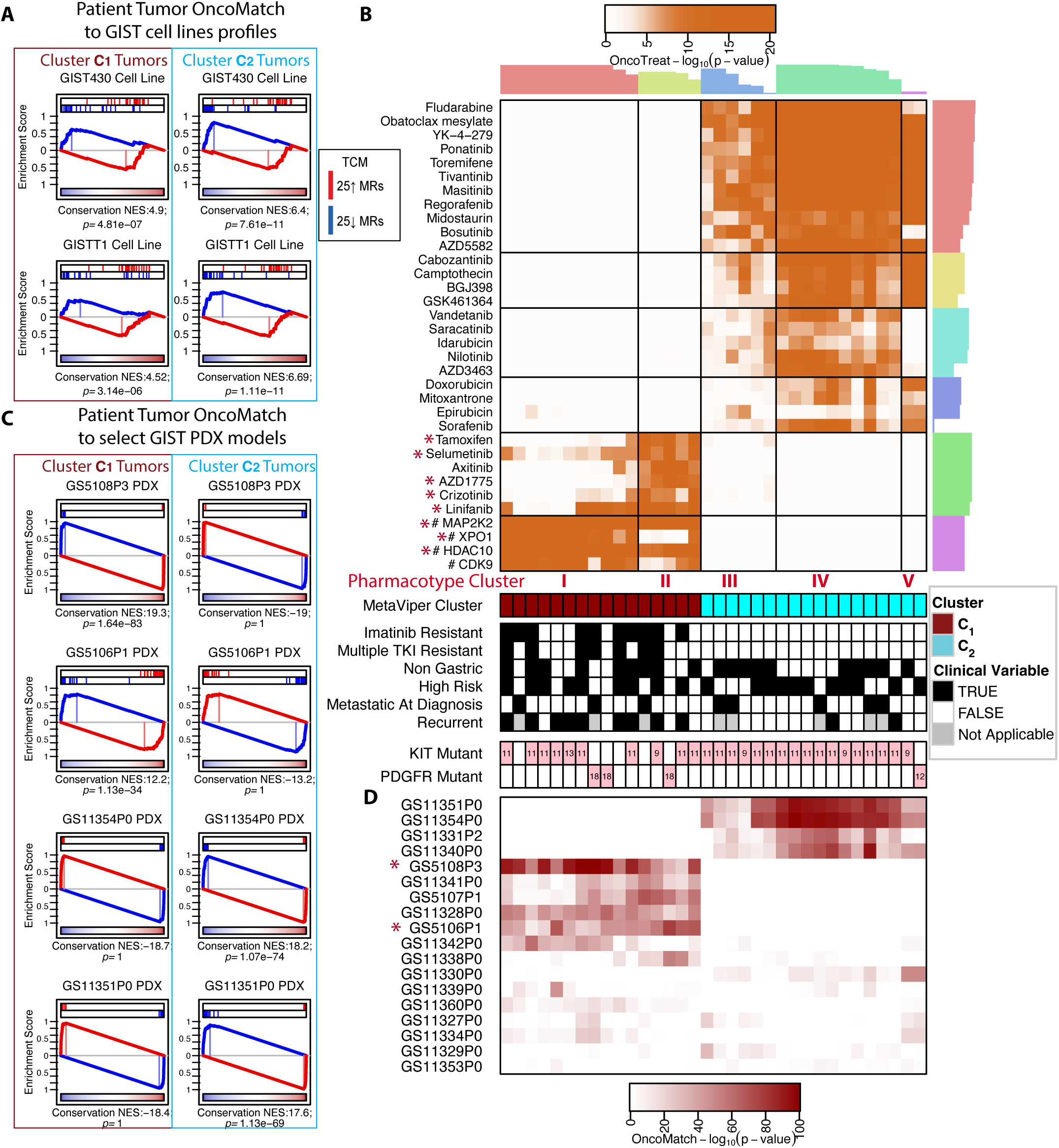
OncoMatch analysis of Life Raft patient tumor samples to available GIST cell line *(**A**)* and patient-derived xenograft (PDX) models *(**C, D**)* and *OncoTreat* drug predictions for patient tumors *(**B**)*. **(A)** RNASeq profiling and protein activity inference using metaVIPER was performed on two available GIST cell lines, GIST-T1 and GIST430, with GIST430 previously reported as less sensitive to imatinib. Integrated protein activity signatures of cluster ***C*_1_** and ***C*_2_** patient tumors were generated. In OncoMatch analysis, we assess the enrichment of the 25 most activated (red bars) and 25 most inactivated (blue bars) patient tumor master regulator (MR) proteins—i.e., the *MR-activity signature*, in the signature of the cell line model. In the plots, proteins are sorted left to right from the most differentially inactivated to the most activated in the cell line model. The NES and associated BH-corrected *p* are displayed. **(B)** *OncoTreat* drug sensitivity predictions for all GIST patient tumors profiled. High throughput perturbation screens with RNASeq profiling using the PLATESeq platform were performed in GIST430 and GIST-T1, with the analysis subsequently focused on GIST430 given its superior match to the imatinib-resistant cluster. *OncoTreat* uses *de novo* drug mechanism information from a carefully selected model (GIST430) to identify MR-activity reversing drugs. For each patient and drug, we compute the enrichment of MRs in the drug signature, as measured by the aREA algorithm, with negative NES indicating reversal, and the associated lower-tail *p*-value. Color intensity in the heatmap corresponds to the −log10 (BH-corrected *p*), and the predictions are clustered by tumor (*columns—*pharmacotype clusters) and drug (*rows*), with cluster reliability indices for cluster assignment shown as barplots on the top and righthand side of the heatmap. A few recurrent *OncoTarget* predictions are incorporated into the heatmap for simplicity and are denoted as [# target protein], with corresponding *OncoTarget* −log10 (BH-corrected *p*). The prior cluster assignment and clinical annotation are provided at the bottom of the heatmap. **(C)** RNASeq and protein activity profiling by metaVIPER of n=18 GIST PDX models developed by Crown Bioscience was performed. The enrichment plots for OncoMatch analysis of the two PDX models most strongly matching to ***C*_1_** (*GS5108* and *GS5106*) and ***C*_2_** (*GS11354* and *GS11351*) patient tumors are shown. **(D)** Heatmap of OncoMatch analysis between individual patient tumors (*columns* organized in same order as *(**B**)*) and all 18 GIST PDX tumors profiled (*rows*), with color intensity corresponding to the −log10 (BH-corrected *p*) for OncoMatch analysis.

We used PLATE-seq^60^ to generate pre- and post-drug perturbation profiles for both GIST-T1 and GIST430, using 46 drugs, including 30 FDA approved antineoplastics and 16 late-stage experimental drugs in phase II or III oncology clinical trials (**Table S5**), following the protocol described in **Methods**. VIPER analysis is used to generate a drug-mediated differential protein activity signature from gene expression profiles of drug vs. vehicle control (DMSO) treated cells. Downstream *OncoTreat* analysis was focused on GIST430 given its superior match to the imatinib-resistant cluster. We have subsequently completed an expanded screen in GIST430, using 333 drugs, including 122 FDA approved antineoplastics, 192 late-stage experimental drugs in phase II or III trials, and 19 compounds from diversity libraries presenting cell line-specific EC_50_ ≤ 2 μM (**Table S6)**. Data from the expanded screen were not yet available for the original *OncoTreat* predictions for *in vivo* validation, but are shared here, including an unsupervised clustering of drug mechanism based on induced differential protein activity (**Figure S4**).

### OncoTreat Predictions

Using data from the GIST430 screen and the *OncoTreat* algorithm, we made tumor-specific predictions for the 34 LRG GIST samples. Samples optimally segregated into five main clusters, based on shared drug predictions (**Figure 3B**). We have previously coined the term *pharmacotype* to describe tumor clusters presenting shared drug sensitivity predictions^28^. Not surprisingly, given the dependency of *OncoTreat* predictions on tumor-specific MRs, all 10 imatinib-resistant tumors aligned into either pharmacotype I or II (**Figure 3B**).

Importantly, we identified five novel candidate drugs, all either FDA approved antineoplastic agents or in late-stage clinical trials, that are predicted to reverse the activity of top MR proteins in subsets of imatinib-resistant GISTs. Specifically, the VEGFR2 and multi-kinase inhibitor linifanib was predicted as significant by *OncoTreat* for 8 of 10 imatinib-resistant tumors (BH-corrected *p* ≤ 10^−5^) and represents the top ranked prediction for four of them. The MEK inhibitor selumetinib was predicted for 5 of 10 imatinib-resistant tumors and is consistent with *OncoTarget* assessed aberrant MAP2K2 activity in ***C_1_***tumors (**Figure 1D**). Finally, the selective estrogen receptor modulator tamoxifen, WEE1 inhibitor AZD1775 (adavosertib), and MET/ALK selective inhibitor crizotinib were each predicted for 3 of 10 imatinib-resistant tumors (BH-corrected *p* ≤ 10^−5^).

In addition to these five *OncoTreat*-predicted drugs, we also considered two *OncoTarget*-based predictions for *in vivo* validation: the selective XPO1 inhibitor selinexor, predicted for 7 of 10 imatinib-resistant tumors, and the pan-HDAC inhibitor panobinostat predicted based on HDAC10 and HDAC7 aberrant activity in 10 of 10 and 7 of 10 imatinib-resistant tumors, respectively (BH-corrected *p* ≤ 10^−5^, by *OncoTarget*, **Figure 1D**). Neither panobinostat nor selinexor were evaluated in our initial GIST430 perturbational screen. The two *OncoTarget* predictions are incorporated into the **Figure 3B** heatmap, for simplicity, and are labeled as [# target protein]. However, the color intensity on the associated rows reports on the aberrant XPO1 and HDAC10 activity (-log10(BH-corrected *p*)).

Perhaps not surprisingly, several of the recurrently predicted drugs for ***C*_2_** tumors (pharmacotypes III, IV, and V) are TKIs with partial selectivity for c-KIT, including masitinib, ponatinib, regorafenib, bosutinib and cabozantinib (**Figure 3B**). Intriguingly, imatinib itself is not predicted by *OncoTreat* for tumors in either cluster. It is possible that our analytical approach, which amplified the signal of MR proteins involved in implementing the imatinib-resistant state, cancels out the signal of MR programs that remain active in both clusters, and perhaps it is these programs that are most efficiently reversed by imatinib.

### Selection of GIST Patient-derived Xenograft Models

Crown Bioscience, Inc, a company that specializes in PDX drug testing, provided us with RNASeq profiles of 18 existing GIST models in their tumor bank. MetaVIPER and OncoMatch analysis (see Methods) identified two high-fidelity models, GS5106 and GS5108, that matched all 10 imatinib-resistant patient tumors (each matched by at least one model at an extremely statistically significant level at p < 10^−30^) (**Figure 3C-D**). Indeed, GS5106 and GS5108 collectively provide highly significant matches to all 16 C_1_ tumors comprising pharmacotypes I and II (columns representing tumors in **Figures 2B** and **2D** are in the same ordering).

Notably, while OncoMatch analysis was completely agnostic to any known clinical or genotypic PDX characteristics, pre-existing therapeutic data—made available by Crown Bioscience and later reproduced in our own studies—confirmed that both models were indeed imatinib resistant. *GS5108* harbors a canonical *KIT* exon 11 mutation and a co-occurring (presumed secondary) *KIT* exon 17 mutation. *GS5106* harbors a *KIT* exon 13 mutation and a co-occurring (presumed secondary) *KIT* exon 17 mutation.

To ensure that the drugs predicted from patient profiles were also predicted for these PDX models, we repeated the *OncoTarget* and *OncoTreat* analyses based on their RNASeq profiles (*GS5106* and *GS5108*), as well as on the profiles of two additional PDX models (*GS11351* and *GS11354*) which were found to represent optimal matches for C_2_ (i.e., not imatinib-resistant) tumors. Of the seven candidate drugs predicted from human tumor profiles, linifanib *[OncoTreat]* and selumetinib *[either by OncoTreat or by OncoTarget based on aberrant MAP2K2 activity]* were predicted as significant in both imatinib-resistant PDX models (**Figure 4A**). In contrast, tamoxifen and AZD1775 *[OncoTreat]* were only predicted as significant in the *GS5106 PDX*, selinexor *[OncoTarget based on aberrant XPO1 activity]* only in the *GS5108*, and crizotinib *[OncoTreat]* was not predicted for either model (**Figure 4A**).

**Figure 4.**
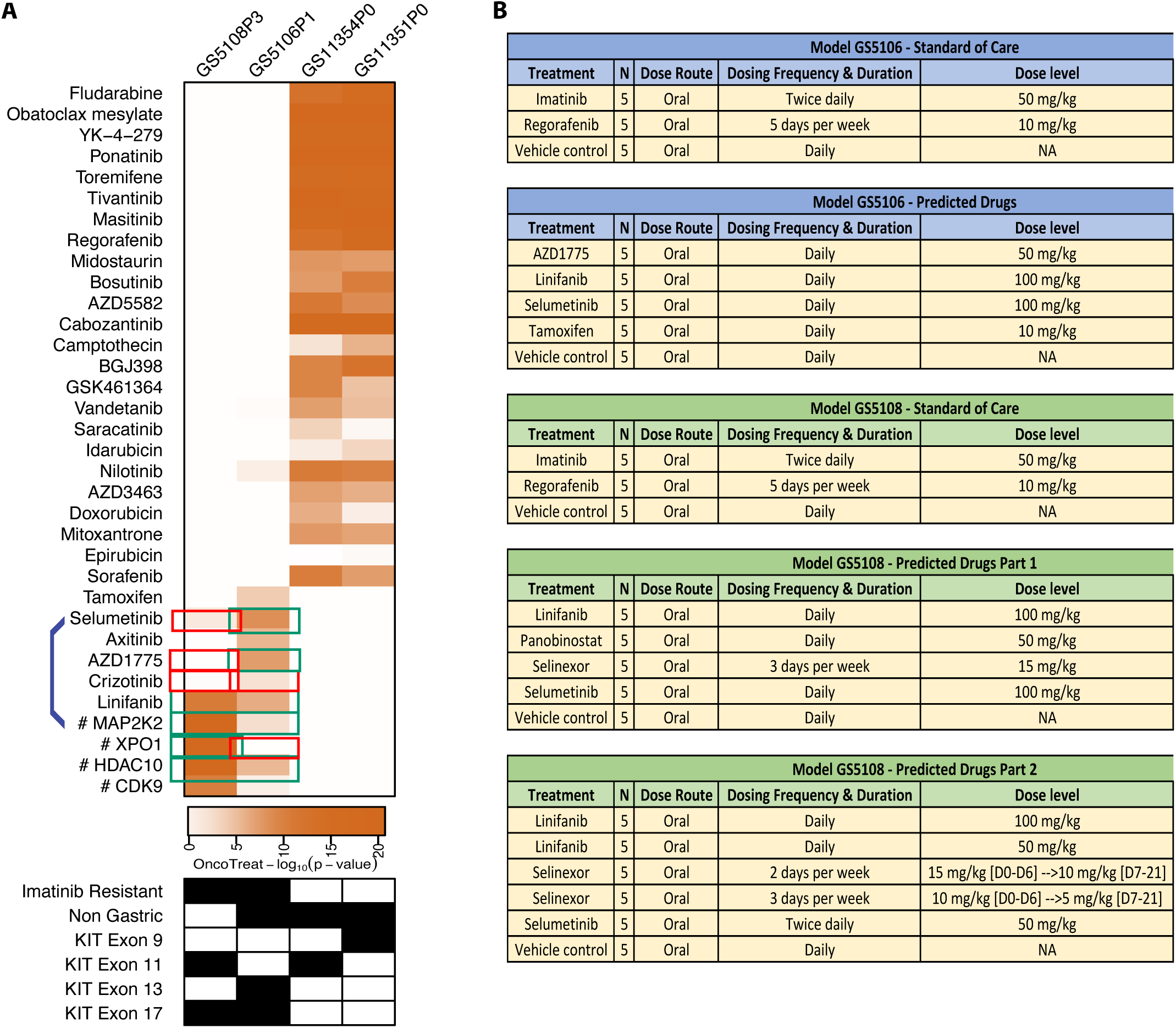
Selection of drug predictions that are concordant in the optimal PDX models and assignment to the therapeutic study. **(A)** Heatmap displays the analogous *OncoTreat* predictions for four GIST PDX models, *GS5108* and *GS5106* that are strong matches to ***C*_1_** (imatinib-resistant) patient tumors and *GS11354* and *GS11351* which are strong matches to ***C*_2_** patient tumors. *Rows* are organized in the same order as in ***Figure 3B***. A few recurrent *OncoTarget* predictions are incorporated into the heatmap for simplicity and are denoted as [# target protein], with corresponding *OncoTarget* −log10 (Bonferroni *p*). Green outlined boxes indicate predictions that were confirmed in the respective PDX, while red outlined boxes indicate that the prediction did not meet statistical significance in the PDX. **(B)** PDX therapeutic study design for *GS5106* and *GS5108*. Only *OncoTreat/OncoTarget* predictions that are concordant for the PDX tumor were incorporated in the study. Initial maximum tolerated dose (MTD) studies in non-tumor bearing NOD/SCID mice were performed at three dose levels for each drug, using reported doses from the literature as an initial starting point. Dose and schedule for all but the final part of the study were based on the MTD.

Notably, assessment of crizotinib as a prediction for the *GS5106* model did not quite meet our statistical significance threshold but was close (*p* ∼ 10^−4^), while it was predicted to have no meaningful activity for *GS5108*. This discordance is consistent with the fact that crizotinib was found to reverse the activity of a subset of tumor-specific MRs detectable in only 3 of 10 imatinib-resistant tumors, and that the activity of this subset of MRs was not strongly conserved in the two PDX models. Thus, crizotinib was eliminated, leaving six candidate drugs for *in vivo* efficacy evaluation. The resulting PDX therapeutic study design is summarized in **Figure 4B**.

The use of metaVIPER was critical to the discovery of a conserved molecular state encompassing imatinib-resistant tumors and in identifying recurrent *OncoTarget* and *OncoTreat*-based drug predictions for these tumors. For comparison, an analogous approach using only differential gene expression signatures (dGES) to perform unsupervised consensus clustering fails to find a stable clustering solution, with k=2 to 4 partitions resulting in trivial solutions with three single-sample clusters, and k=5 (**Figure S5A**) resulting in two larger but ambiguous groupings that do not cleanly separate imatinib-resistant tumors. Likewise, the use of dGES to select druggable targets based on high mRNA expression (**Figure S5B**) and to select drugs that reverse tumor-specific dGES of imatinib-resistant tumors (**Figure S5C**) fails to identify recurring drug predictions for these tumors. The overexpression of *MAPK7* has been reported as predictive of linifanib response in GIST cell lines^61^, and indeed is overexpressed in the group of ***C_1_***vs. ***C_2_*** tumors in our dataset (*p*=6.51e-06, by Wilcoxon rank-sum), but the differential mRNA Z-score is ≥ 2.0 for only one of 16 ***C_1_*** tumors, limiting its predictive ability on an individual tumor basis.

### OncoTreat and OncoTarget Predict Treatment Response in PDX Models

*In vivo* studies proceeded in four phases: 1) maximum tolerated dose (MTD) finding studies in non-tumor bearing NOD/SCID mice; 2) therapeutic studies with standard of care drugs for GIST (imatinib [negative control] and regorafenib) in P3 passages of *GS5106* and *GS5108*; 3) therapeutic studies with *OncoTreat* or *OncoTarget*-predicted drugs; 4) testing of active predicted drugs at titrated doses lower than the MTD. Early on-treatment biopsy specimens were collected in each of the therapeutic arms (in phases 2-4), for pharmacodynamic testing; these mice were excluded from therapeutic response assessment.

MTDs identified were: imatinib 50 mg/kg twice daily; regorafenib 10 mg/kg daily (5 days on, 2 days off); linifanib 100 mg/kg daily; selumetinib 100 mg/kg daily; tamoxifen 10 mg/kg daily; AZD1775 50 mg/kg daily; panobinostat 50 mg/kg daily; and selinexor 15 mg/kg three times weekly. Predicted drugs were tested only in the PDX model(s) in which they were also predicted, as summarized in **Figure 4B**. Mice were treated for 21 days, with tumor measurements continuing for 28 days. Treatment doses were held as needed based on total body weight loss parameters, per protocol, and other observed toxicities.

Therapeutic studies in both PDX models confirmed complete phenotypic resistance to imatinib with no significant difference in tumor growth between imatinib and vehicle control-treated mice (**Figure 5A&C**). This is reassuring as these models were selected agnostic to any known clinical, genotypic, or phenotypic characteristics of the PDX, but rather based on their ability to recapitulate the MR signatures we saw in imatinib-resistant patient tumors. Importantly, the multi-kinase inhibitor regorafenib demonstrated significant mean tumor growth inhibition (TGI) on treatment relative to vehicle control in both models, achieving 55% TGI in *GS5106* (*p* = 0.022 based on a linear mixed model for repeated measurements as implemented in^62^) and 120% TGI in *GS5108* (i.e., 20% regression from baseline, *p* = 0.002) (**Figure 5A&C**). While regorafenib is clinically approved for imatinib-resistant GIST, it is often poorly tolerated for long term use in patients due to multiple off-target side effects^63^.

**Figure 5.**
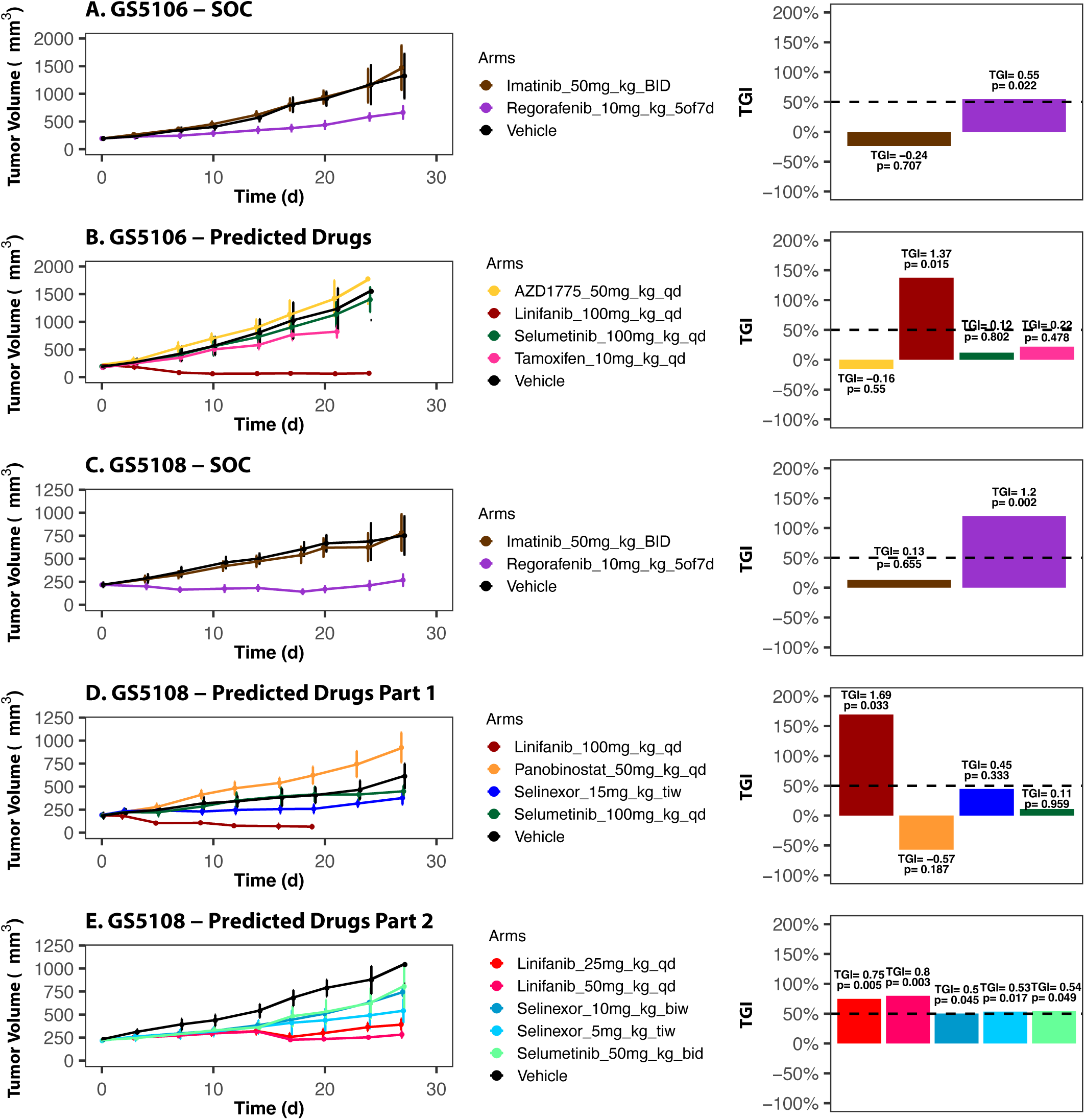
Treatment response in GIST PDX models. Therapeutic PDX studies were conducted in 5 parts (***A-E***) in collaboration with Crown Bioscience, each with a vehicle control arm. **(A)** Standard of care drugs imatinib and regorafenib were tested in *GS5106*. Tumor growth curves with standard error estimates (left panel) and relative tumor growth inhibition (TGI) on treatment compared to vehicle control (right panel) are shown. The percentage of TGI relative to vehicle control and associated linear mixed model *p*-value for TGI are displayed for each arm in the barplot. **(B)** Drugs predicted by *OncoTreat* (AZD1775, linifanib, selumetinib, and tamoxifen) were tested in *GS5106*. **(C)** Standard of care drugs imatinib and regorafenib were tested in *GS5108*. **(D)** Drugs predicted by *OncoTreat* (linifanib, selumetinib) and *OncoTarget* (selinexor based on XPO1 activation and panobinostat based on HDAC10/7 activation) were tested at the maximum tolerated dose in *GS5108*. Panobinostat resulted in accelerated tumor growth compared to vehicle control. **(E)** Because part 4 of the study (*panel D*) suffered from unexpected toxicity of unclear etiology, resulting in weight loss and early death in mice across multiple arms, which prompted significant dose interruptions and dropout of mice, an additional study was performed evaluating linifanib, selinexor, and selumetinib at multiple dose levels. Linifanib continued to demonstrate significant TGI across dose levels. Likewise, both selinexor arms and the selumetinib arm demonstrated statistically significant TGI in the 50-55% range, at the lower dose levels, but with more consistent dosing.

Linifanib, which was predicted by *OncoTreat* for 8 of 10 imatinib-resistant LRG tumors, resulted in marked tumor regression in both PDX models, with 137% TGI (*p* = 0.015) and 169% TGI (*p* = 0.033) in *GS5106* and *GS5108*, respectively, when evaluated at the MTD (**Figure 5B&D**). Selinexor, which was predicted by *OncoTarget* for 7 of 10 imatinib-resistant tumors based on XPO1 activation, was evaluated in *GS5108* and resulted in 45% TGI on treatment (**Figure 5D**). This is close to the commonly cited TGI ≥ 50% benchmark in PDX therapeutic studies^64–66^ that predicts clinical benefit. Selumetinib, which was predicted for 5 of 10 imatinib-resistant patient tumors and in both PDX models by *OncoTreat*, unfortunately did not result in significant TGI in either *GS5106* or *GS5108* (**Figure 5B&D**). Likewise, tamoxifen and AZD1775, which were predicted for a smaller subset of imatinib-resistant tumors and only tested in *GS5106*, demonstrated no significant TGI (**Figure 5B**). Finally, panobinostat was evaluated in *GS5108* and demonstrated no significant TGI and in fact apparently resulted in acceleration of tumor growth with −57% TGI (**Figure 5D**).

There were concerns with the health of the cohort of mice used for these therapeutic studies, including premature deaths across multiple arms, despite all drug doses being well tolerated in the prior MTD studies in non-tumor bearing mice. Animal-level data across study duration for tumor volume, medication administration record, and body weight are provided in **Table S7**, with deaths highlighted in red. As these studies were being conducted in the early days of the COVID-19 pandemic, it is possible that sterility conditions in the testing facility were suboptimal. The dropout of mice in the therapeutic study further limited statistical power to confirm meaningful TGI—e.g., the 45% TGI with selinexor in *GS5108* did not meet statistical significance due to fewer mice making it to the end of treatment (EoT) timepoint. We thus proceeded to perform a final phase of therapeutic testing, evaluating the three strongest predictions in *GS5108*, linifanib, selumetinib, and selinexor, at multiple dose levels.

The efficacy of linifanib at inducing significant TGI was demonstrated at doses significantly lower than the MTD, 50 mg/kg and 25 mg/kg daily, with TGI of 80% (*p* = 0.003) and 75% (*p* = 0.005), respectively, although there was some clear dose dependence on the extent of response (**Figure 5E**). There was also moderate TGI with selinexor, which after a one-week run-in at the MTD, was administered at the lower tested doses, 10 mg/kg twice weekly and 5 mg/kg three times weekly, with TGI of 50% (*p* = 0.045) and 53% (*p* = 0.017), respectively. Selumetinib also resulted in moderate TGI of 54% (*p* = 0.049) at the alternative dosing schedule, 50 mg/kg twice daily, in *GS5108* (**Figure 5E**). It is likely that the greater benefit seen with selumetinib at the lower dose was due to the ability to more consistently administer the dose, as the mice were generally healthier in this final phase of testing.

A comprehensive summary of TGI assessment for all therapeutic arms and statistical significance testing is presented in **Table S8**.

### Pharmacodynamic Assessment of MR-activity reversal by Predicted Drugs *In Vivo*

Pharmacodynamic (PD) studies are critical to drug development to elucidate mechanism and to characterize resistance. Specifically, we were interested in learning if *OncoTreat*-predicted drugs that demonstrate efficacy in the PDX, recapitulate *in vivo* the reversal of MR activity that occurs in the cell line model and forms the basis of prediction. Additionally, since a number of our predictions were ineffective, our PD analysis attempted to determine if these drugs were ineffective due to an inability to achieve MR-activity reversal *in vivo*, perhaps due to pharmacokinetic factors, or ineffective despite achieving reversal, for instance due to later cell adaptation. Finally, as regorafenib, one of the standard of care arms, demonstrated effective TGI in the PDXs despite not being predicted for imatinib-resistant tumors by *OncoTreat*, we wished to learn if regorafenib induced MR-activity reversal *in vivo* in these two models.

Biopsy specimens were collected and analyzed by VIPER to assess MR-activity reversal in drug vs. vehicle control-treated PDX samples, **see Methods**. In *GS5106*, ineffective drugs including imatinib and the predicted drugs AZD1775, tamoxifen, and selumetinib, demonstrated no statistically significant evidence of *in vivo* MR-activity reversal (**Figure 6A**). On the other hand, both effective drugs, including the standard of care control regorafenib (*p* = 9 x 10^−17^) and linifanib (*p* = 2 x 10^−18^, by *OncoTreat*) demonstrate significant MR-activity reversal.

**Figure 6.**
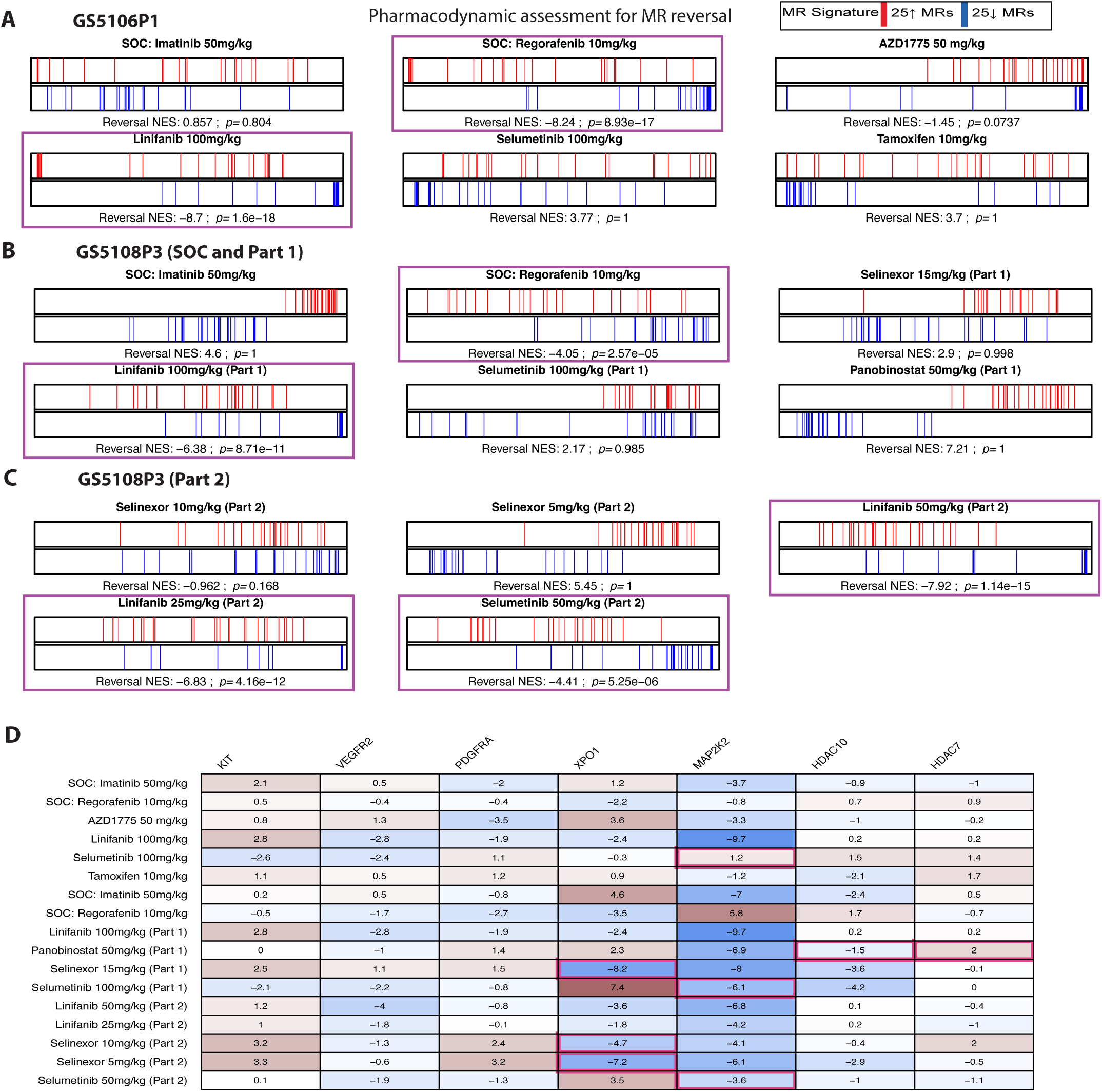
Pharmacodynamic assessment of MR-activity reversal *in vivo* and activity changes in selected *OncoTarget* proteins. In the two GIST PDX models, two mice from each drug arm were sacrificed after the 3^rd^ dose. VIPER was used to generate a differential protein activity signature for each drug-treated versus corresponding vehicle control-treated arm. Reversal of the MR-activity signature, as defined from the baseline PDX profiles, was assessed in the treatment signature by enrichment analysis (aREA) *(**A-C**)*. Statistically significant MR-activity reversal *in vivo* (BH-corrected *p* < 10^−5^, highlighted by surrounding purple boxes) was confirmed for linifanib in both PDX models, including all three dose level arms in *GS5108*. Regorafenib, an approved clinical option for GIST after progression on imatinib but not predicted for ***C_1_*** tumors in our analysis, also demonstrated significant MR-activity reversal in both models, consistent with its significant TGI *in vivo*. Selumetinib, which resulted in TGI in *GS5108* at the lower 50 mg/kg dose, demonstrated significant MR-activity reversal only at this dose level. Drug predictions by *OncoTreat* that did not induce significant TGI generally did not induce significant MR-activity reversal *in vivo*, including the AZD1775, selumetinib, and tamoxifen arms in *GS5106*. **(D)** Changes in activity of select druggable proteins are shown in the heatmap. Boxes outlined in pink link OncoTarget predicted drugs and their canonical targets, e.g., Selinexor -- XPO1 and Panobinostat -- HDAC10/HDAC7.

Likewise, in *GS5108*, imatinib demonstrated no significant *in vivo* MR-activity reversal (**Figure 6B**). On the other hand, standard of care control regorafenib demonstrated borderline MR-activity reversal (*p* = 3 x 10^−5^) and linifanib demonstrated marked reversal across multiple dose levels (*p* = 9 x 10^−11^ at 100 mg/kg daily, *p* = 1 x 10^−15^ at 50 mg/kg daily, and *p* = 4 x 10^−12^ at 25 mg/kg, all by *OncoTreat*) (**Figure 6B-C**). Interestingly, selumetinib, which demonstrated moderate TGI at the lower and better tolerated dose level, demonstrated MR-activity reversal at this lower 50 mg/kg dose (*p* = 5 x 10^−6^, by *OncoTreat*) but not at the 100 mg/kg dose (**Figure 6B-C**). The effect seen in each drug arm on individual top candidate MRs (25 most activated and 25 most inactivated) of the respective PDX model, is presented in **Figure S6**.

Selinexor, which also demonstrated significant TGI compared to control, did not achieve *in vivo* MR-activity reversal (**Figure 6B-C**). However, selinexor does markedly decrease XPO1 activity, which formed the basis of its prediction by *OncoTarget*, *in vivo*; conversely, panobinostat, which was ineffective, did not significantly reduce HDAC10/7 activity (**Figure 6D**). In fact, panobinostat is noted to significantly enhance MR-activity in *GS5108* (**Figure 6B**), corresponding to the accelerated tumor growth we observed. Interestingly, selumetinib was the only drug that demonstrated a trend toward reduction in c-KIT activity in these PDX models.

Overall, there was modest anticorrelation between TGI% and MR-activity reversal across all treatments arms, Spearman-rank correlation ρ = −0.72, associated p-value = 0.001567. In treatments arms specifically predicted by OncoTreat, the ρ =-0.75, p-value = 0.02549.

## Discussion

The development of high-affinity inhibitors of the c-KIT tyrosine kinase, most notably imatinib^67^, has greatly benefited patients with GIST, a disease characterized by aberrant c-KIT activity^3^. C-KIT targeting therapy has reduced the rate of relapse after surgical resection of high-risk GIST^68^ and significantly prolonged survival in the metastatic setting^12,25^. Unfortunately, the eventual development of imatinib resistance is common, and is associated with poor outcomes^69^. While most tumors that develop acquired resistance to imatinib harbor secondary mutations in *KIT*^16,17^, next-generation c-KIT inhibitors like ripretinib designed, to overcome these mutations, have achieved objective response rates of 9% and mPFS of 6.3 months in the imatinib-resistant setting^26^.

In this study, we applied *OncoTarget* and *OncoTreat—*RNA-based methodologies that identify candidate drugs that target Master Regulator (MR) proteins—to the problem of imatinib-resistant GIST. These approaches are designed to complement traditional oncogene-centered precision cancer medicine by identifying dependencies arising from the broader molecular landscape of a tumor. Oncogene mutations represent only one of many potential mechanisms for inducing the aberrant activity of a protein^37^. Conversely, the presence of a mutation does not guarantee drug sensitivity, as secondary mutations can drive selection of resistant cells^38^.

*OncoTarget* identifies individual MRs with available high-affinity inhibitors, while *OncoTreat* identifies drugs capable of reversing the activity of the 25 most aberrantly active and 25 most aberrantly inactive MRs of a tumor—a module dubbed the *Tumor Checkpoint* (TCM)^37^ and shown to be responsible for canalizing the effect of >80% of functional mutations in individual tumors^70^. *OncoTreat* does so by leveraging large-scale drug perturbation screens with RNASeq profiling of the molecular response in carefully selected *in vitro* models to identify clinically relevant drugs that reverse MR activity. As shown here, *OncoTarget* and *OncoTreat* provide scalable tools to efficiently prioritize candidate therapies for drug-resistant tumors.

While *OncoTarget* and *OncoTreat* predictions have been systematically tested in PDX models^28^, a distinguishing feature of the present study is their application to tumors that fail to respond or relapse following standard of care therapy. Imatinib resistance is a major determinant of poor outcome in advanced GIST.

Our analysis shows that MR protein activity is markedly distinct in imatinib-resistant vs. imatinib-sensitive tumors (**Figure 1**), with substantial conservation of aberrantly activated and inactivated proteins across imatinib-resistant GISTs, independent of whether imatinib resistance was primary or acquired. This conservation suggests that imatinib resistance is not fully accounted for through the clonal selection of secondary mutations that reduce imatinib binding and restore c-KIT activity, but rather also involves a molecular reprogramming of cell state. Indeed, analysis of scRNA profiles of high-risk GIST demonstrates evidence of treatment-emergent subpopulations strongly enriched in this same MR-activity signature, with the proportional expansion of these subpopulations associated with clinical progression on imatinib (**Figure 2**). Furthermore, subpopulations strongly enriched in this MR signature demonstrate modestly decreased c-KIT activity and no association with *KIT* mRNA expression or new inferred copy number alterations. Notably, six tumors that had not yet developed overt clinical progression on imatinib present substantial MR activity overlap with those presenting frank imatinib resistance, three of which would also be predicted to be imatinib-resistant based on genotype (two being *KIT*/*PDGFRA*-wildtype and one harboring a *KIT* exon 13 primary mutation associated with relative resistance, **Figure 1**). Whether these tumors will ultimately develop clinical progression on imatinib requires further follow up—although some may have been cured by timely surgical resection alone.

Equally important, we show that the VIPER-based OncoMatch algorithm—originally developed to identify high-fidelity models of human tumors that recapitulate their MR activity^29^— could identify only a small subset of 18 established GIST PDX models as optimally recapitulating the imatinib-resistant cluster MR signature. This has two important implications. First, it allows effective validation of predicted drugs in models that most closely capture the clinical drug-resistant state. Second, and perhaps more importantly, it avoids testing drugs in models that fail to replicate the activity of imatinib-resistant GIST, which could lead to misleading results.

There were several limitations to our study. First, as expected, not all predictions were effective. The PD studies show that several ineffective *OncoTreat-*predicted drugs also failed to achieve MR-activity reversal *in vivo* (**Figure 6**), suggesting that pharmacokinetic barriers may have contributed to their lack of efficacy, since in a previous study, 15 of 18 *OncoTreat*-predicted drugs induced MR-activity signature reversal *in vivo* and resulted in disease control. Second, the drug doses evaluated in our therapeutic studies, determined from MTD studies in non-tumor bearing mice, were not always consistent with previously published PDX studies (e.g., we tested regorafenib at a dose of 10 mg/kg 5 days on, 2 days off, whereas in other studies it has been tested at doses as high as 20 mg/kg daily^71^), which may impact the translatability of our findings. Dosing differences may reflect mouse strain-specific differences in pharmacokinetics and the inherent variability of mouse MTD studies, which rely on limited endpoints—i.e., body weight and death. Third, there is not an optimal negative control to evaluate our predictions against. Because MR-activity reversal is only one of several potential mechanisms of anti-tumor response, we have not established the negative predictive value of *OncoTarget* or *OncoTreat*. In a previous study across several cancer types, a panel of 13 randomly selected unpredicted drugs served as negative controls and proved ineffective^28^; here, imatinib served as the sole negative control.

Subsequent to the original validation studies, we have completed a larger perturbational screen in GIST430, incorporating 333 clinically relevant oncology drugs. These new profiles have helped identify several additional candidate drugs by *OncoTreat* that are predicted—but not yet validated—to induce MR-activity reversal in a sizable subset of imatinib-resistant tumors (**Figure 7**: pharmacotypes III & IV). Among others, these include axitinib, MLN-2480 (pan-RAF inhibitor), AZD-5069 (CXCR2 antagonist), savolitinib (MET inhibitor), prinomastat (matrix metalloproteinase inhibitor), ipatasertib (pan-AKT inhibitor), abexinostat (pan-HDAC inhibitor), enasidenib (IDH2 inhibitor), conventional alkylating chemotherapeutics [carmustine and mechlorethamine], multiple hormonal signaling modulators [enzalutamide, fulvestrant], and differentiating agents [ATRA (vitamin A metabolite) and arsenic trioxide]. Whether combinations of MR-targeting agents with imatinib or other TKIs that directly target c-KIT could achieve greater efficacy remains an open question, particularly given evidence that imatinib-sensitive and resistant cell states may co-exist within individual tumors^72^, albeit at different ratios (**Figure 2**). The recently completed phase III PEAK trial in imatinib-resistant GIST demonstrated a significant benefit in PFS when the standard of care drug sunitinib—which like linifanib is a VEGFR2 and multi-kinase inhibitor—is combined with the next-generation c-kit inhibitor bezuclastinib, compared to sunitinib alone, with mPFS 16.5 vs. 9.2 months (HR 0.50)^73^, establishing the feasibility and potential benefit of such combinations.

**Figure 7.**
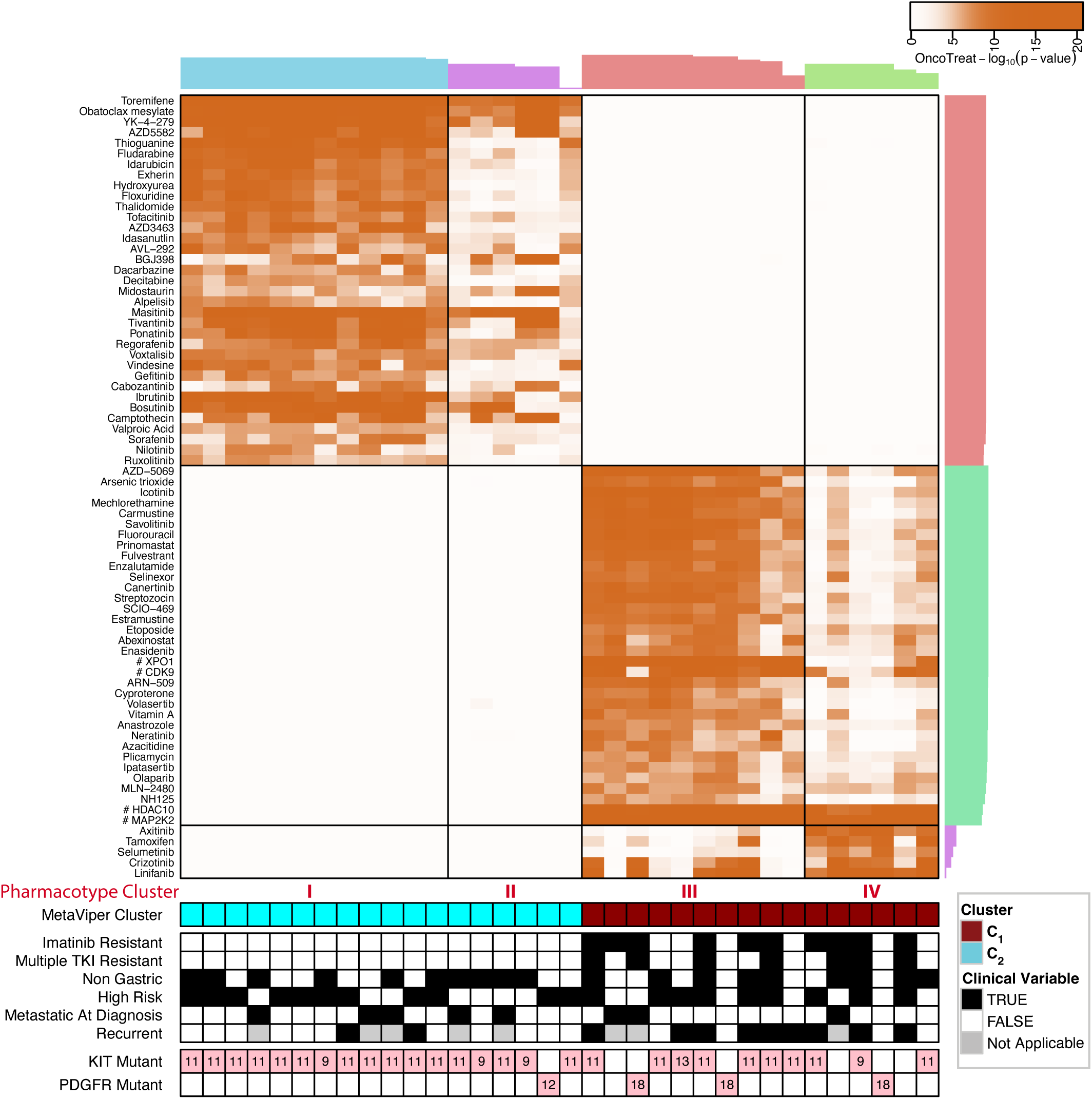
*OncoTreat* predictions for LRG patient tumors after completion of an expanded high throughput drug screen in GIST430, using 333 drugs, including 122 FDA approved antineoplastics and 192 late-stage experimental drugs in phase II or III trials. (**A**) Heatmap of *OncoTreat* predictions using data from the expanded drug screen in GIST430. Color intensity in the heatmap corresponds to the −log10 (BH-corrected *p*), and the predictions are clustered into pharmacotypes (*columns*) and by drug (*rows*), with cluster reliability indices for cluster assignment shown as barplots on the top and righthand side of the heatmap. A few recurrent *OncoTarget* predictions are incorporated into the heatmap for simplicity and are denoted as [# target protein], with corresponding *OncoTarget* −log10 (BH-corrected *p*). The prior cluster assignment and clinical annotation are provided at the bottom of the heatmap. Several additional candidate drugs to target imatinib-resistant tumors emerge. Interestingly, while panobinostat is not predicted based on its measured effect in GIST430, an alternative pan-HDAC inhibitor abexinostat is strongly predicted for several imatinib-resistant tumors. Selinexor is also predicted by *OncoTreat* for the vast majority of tumors with XPO1 activation.

The framework discussed here may be applicable to other cancer types in which drug-resistant states are associated with conserved transcriptional reprogramming. The approach could also be applied iteratively as resistance evolves over time, though this requires longitudinal sampling and will benefit from advances in blood-based tumor RNA profiling^74^. Importantly, we have demonstrated the value of VIPER-based protein activity inference applied to scRNASeq profiles^75,76^, and *OncoTreat* and *OncoTarget* can in principle be applied to optimally target dominant subpopulations within tumors. Targeting of multiple subpopulations, either sequentially or in combination, may ultimately be necessary to durably overcome resistance.

## Methods

### Generation of Gene Regulatory Networks

To support context-specific regulatory protein activity inference by VIPER and metaVIPER, we have generated comprehensive molecular interaction networks (interactomes) using the Algorithm for the Reconstruction of Accurate Cellular Networks (ARACNe)^77,78^. ARACNe requires ≥ 100 samples to produce accurate predictions. Since GIST repositories are much smaller, we leveraged metaVIPER, an extension of VIPER that effectively infers a protein’s activity by integrating information on its transcriptional targets from multiple regulatory networks. In this case, we used metaVIPER to integrate regulatory networks representing the cellular context of 43 distinct cancer types or subtypes^45,46^.

Most networks were reverse engineered through the application of ARACNe to cohorts from The Cancer Genome Atlas (TCGA), whenever ≥ 100 RNASeq profiles were available. While there is a soft tissue sarcoma cohort in TCGA (“sarc”), GISTs were specifically excluded. The TCGA RNASeq level 3 data were downloaded from NCI Genomics Data Commons^79^. Raw counts were normalized and variance stabilized by fitting the dispersion to a negative-binomial distribution as implemented in the DESeq2 R-package^80^ (RRID: SCR_015687).

ARACNe was run with 100 bootstrap iterations using an input set of candidate regulators including: (a) 1,810 transcription factors annotated in the Gene Ontology (GO)^81^ “Molecular Function database” as DNA-binding transcription factor activity (GO:0003700), DNA binding (GO:0003677), Transcription regulator activity (GO:0030528), or Regulation of transcription, DNA-templated (GO:0003677 and GO:0045449); (b) 668 transcriptional co-factors manually curated from genes annotated as Transcription Coregulator Activity (GO:0003712), Plays a Role in Regulating Transcription (GO:0030528), or Regulation of Transcription (GO:0045449); (c) 3,483 genes encoding for signal transduction proteins, dually annotated in the GO “Biological Process database” as Signal Transduction (GO:0007165) and in the GO “Cellular Component database” as either Intracellular (GO:0005622) or Plasma membrane (GO:0005886); and (d) an additional 1,905 genes encoding for cell surface proteins (GO:0009986) not categorized as signaling proteins. The Data Processing Inequality (DPI) parameter of ARACNe was set to 0, the Mutual Information (MI) threshold set to *p* = 10^−8^, and the mode of regulation computed based on the correlation between regulator and target gene expression as previously described^32^. The version of n=43 generated networks used in our work is provided (see Figshare: https://figshare.com/articles/dataset/Available_Interactomes_Networks_/30246550).

### Collaboration with Life Raft Group

We have worked closely with the patient advocacy organization, Life Raft Group (LRG), the world’s largest private sponsor of GIST research, which agreed to share with us coded (deidentified) tumor bank specimens and linked clinical annotation. Available clinical variables included primary tumor location, stage at diagnosis, risk classification at diagnosis based on tumor size and mitotic rate, clinical genotype testing (specific *KIT* and *PDGFRA* variants), subsequent disease status (relapse/metastasis), survival from diagnosis, and prior treatments and durations.

### Nucleic acid extraction, sequencing, and analysis

All formalin-fixed paraffin-embedded (FFPE) samples received from LRG were sectioned into fresh slides and stained by H&E, evaluated by a pathologist, and micro-dissected to ensure that a minimum of 70-80% viable tumor was present for subsequent extraction and analyses. RNA was extracted using the QIAGEN RNeasy (FFPE) kit (RRID: SCR_008539). Strand specific sequencing libraries were generated from rRNA depleted total RNAs using the NEBNext Ultra Directional RNA Library Prep Kit for Illumina. Sequencing was performed on the Illumina NextSeq 500 sequencer (RRID:SCR_014983) with 100bp x 2 paired-end reads to a depth of 20M. We initially performed RNASeq on about 50 samples from the LRG tumor bank, including several imatinib-resistant cases. Some samples did not meet quality control standards following library generation or sequencing, and thus the final analysis dataset included 34 unique tumors, including 10 that had clinical progression on imatinib. Alignment was performed with STARv2.5.3^82^ (RRID:SCR_004463) to the Genome Reference Consortium GRCh38.v90 build with gene-level raw counts summarization with STAR quant-mode.

### metaVIPER Analysis

We have previously extensively validated the Virtual Proteomics by Enriched Regulon (VIPER) algorithm as a highly robust and specific tool for the accurate inference of regulatory protein activity in a tissue context-dependent manner^32,37,42^. VIPER leverages accurate tissue-specific gene regulatory networks to measure differential protein activity from bulk or single-cell gene expression signatures (GES). Specifically, akin to a multiplexed gene-reporter assay, VIPER measures a protein’s differential transcriptional activity through a probabilistic enrichment framework that assesses the normalized enrichment score (NES) of its activated and repressed transcriptional targets (*regulon*) in genes over and under expressed in a sample of interest compared to a set of control samples (*reference model*).

Given significant batch effects introduced through various mRNA enrichment strategies for library generation from FFPE samples, the only suitable *reference model* for our analysis of LRG GIST tumors was the centroid of all gene expression profiles comprising the dataset. Thus, we computed a differential gene expression signature (dGES) for each tumor sample as the Z-score of each gene’s expression relative to its distribution across all 34 LRG samples. Next, VIPER, using each regulatory protein’s set of regulated genes, as assigned in the network, performs an enrichment test to determine if the protein’s transcriptional footprint is over-, under-, or normally represented in the tumor’s dGES. VIPER assigns each protein this normalized enrichment score (NES) as a measure of its regulatory activity. Specifically, VIPER implements a rapid analytic-rank based enrichment analysis (aREA) test^32^, which efficiently provides test statistics similar to permutation-based gene set enrichment analysis^52^. When a suitable context-specific gene regulatory network is not available, as was the case for GIST, we have successfully employed metaVIPER, which essentially first runs VIPER using all available networks independently, and then for each protein performs Stouffer’s Z-score integration for each of the independently derived NES’s, which can optionally be weighted^46^.

The most differentially active proteins identified by VIPER comprise candidate Master Regulator (MR) proteins that mechanistically control a tumor’s transcriptional identity, as shown by multiple studies, see^37^ for a comprehensive perspective. VIPER reproducibility is extremely high, such that Spearman’s rank correlation of activity profiles generated from RNASeq at 30M to as low as 50K read depth^32^ is p > 0.8, even though correlation of the underlying gene expression profiles is low p < 0.3. While VIPER is uniquely suited for assessing regulatory proteins that directly control gene expression, including transcription factors, co-factors, and chromatin remodeling enzymes, we have shown that the algorithm is equally effective in monitoring activity of signaling proteins^32^ and cell surface markers^46^.

### Optimal Clustering Analysis

Consensus and optimal clustering were performed on the metaVIPER derived differential protein activity profiles of the 34 LRG GIST tumors, as implemented in the ConsensusClusterPlus^48^ (RRID:SCR_016954) and OptCluster^47^ packages for R. Specifically, a partition around medoids (PAM) approach, using Spearman-rank correlation as the distance metric between samples, considering k=2 to 8 clusters (8 representing 0.25 * the total number of samples), and performing n=10,000 iterations was employed for the analysis presented in **Figure 1**. All relevant statistical analyses were performed using R software (v4.5.2) (RRID:SCR_001905). For comparison, the same consensus and optimal clustering approach was performed using differential gene expression signatures (mRNA Z-scores) relative to the centroid, instead of protein activity as its input, and is presented in Figure S5.

As the optimal clustering solution grouped samples into two roughly even partitions, metaVIPER was run iteratively, this time using the centroid of samples from the opposing cluster (instead of the entire dataset) to generate the dGES, in order to further amplify the protein activity signal derived from the imatinib-resistant (cluster ***C*_1_**) state. The obtained protein activity signatures were used for further downstream analysis, including *OncoTarget* and *OncoTreat* predictions.

### Pathway Enrichment Analysis

Pathway analysis on the integrated differential protein activity signature of cluster ***C_1_*** [imatinib-resistant] vs. ***C_2_*** tumors, as presented in supplementary **Figure S1**, was performed with ‘Gene Ontology’, ‘Cancer Hallmarks’, ‘Immunological Signatures’, and ‘Oncogenic Signatures’ gene sets provided in the Broad MSigDB collections^51–54^ (RRID:SCR_016863). Pathway enrichment analysis was performed by using the single-tail aREA method described above as a rapid approximation to the Kolmogorov-Smirnov (KS)-test used in GSEA^52^, inputting the pathway genes and the sorted differential protein activity signature. Unlike other statistical tests used for pathway enrichment analysis, such as the Fisher’s exact test on a thresholded list of differentially expressed genes/proteins or the repeated application of the KS test in classical GSEA, aREA does not require the arbitrary binarization into significant/non-significant hits for target genes in the signature but takes into account their relative position in the signature.

### Analysis of scRNA Seq Dataset

Raw SRA files were downloaded from gene expression omnibus (GEO: Series GSE254762) and converted to fastq format. As libraries were generated using Singleron’s GEXSCOPE 3’-end sequencing kit^44,83^, the CeleScope pipeline (RRID:SCR_023553) was used with its default parameters for integrated read alignment to GRCh38.v110, unique molecular identifier (UMI) filtering and deduplication, and cell-calling. Cells were further filtered for quality by removing cells with mitochondrial RNA exceeding > 25% and cells with fewer than 500 or greater than 4,000 mapped features, using the Seurat package for R (SCR_016341)^84^. Next, SingleR^85^ (RRID:SCR_023120) was used to annotate all cells with a cell type represented in the Human Primary Cell Atlas^86^.

The malignant cell compartment was then isolated based on (1) expression of *ANO1* (also known as *DOG1*, a highly specific GIST marker used in histopathological diagnosis^87^), (2) SingleR^85^ annotation of cell type (restricted to smooth muscle cells, fibroblasts, mesenchymal stem cells, tissue stem cells, chondrocytes, neuroepithelial cells and neurons), and (3) further refined based on inference of global copy number variation^88^. InferCNV (RRID:SCR_021140) was used to identify inferred copy number variations, using cells annotated as immune system cells by SingleR^85^ (T cells, NK cells, Monocytes, and Macrophages) as reference. A total copy number score was computed for each candidate malignant cell based on the gene-specific absolute values outputted by InferCNV, and the distribution of scores for each sample were fitted to a bimodal gaussian distribution to define a lower cut-off for removal. The plots in Figure S3 are generated using the pheatmap package (RRID:SCR_016418) for R instead of InferCNV’s built-in plotting function.

Finally, raw counts for the malignant cell compartment were normalized and variance stabilized as implemented in the DESeq2 R-package^80^, dGES’s were computed for each cell relative to the gene-wise centroid of malignant cells across all 9 tumor samples, and metaVIPER was employed to infer differential protein activity. Uniform Manifold Approximation and Projection (UMAP projections) for each sample using the first 30 principal components of the matrix of protein activity signatures were generated using the umap package for R (RRID:SCR_018217). The aREA algorithm was again used to evaluate for the enrichment of the integrated ***C_1_*** imatinib-resistant cluster signature (ImResS) in the protein activity signature of each malignant cell.

### OncoTarget Analysis

Through the use of (a) DrugBank^78^ (RRID:SCR_002700), (b) the SelleckChem database (RRID:SCR_003823)^89^, (c) published literature, and (d) publicly available information on pharmaceutical company drug development pipelines, we have curated a refined list of 180 *actionable proteins* representing validated targets of high-affinity pharmacological inhibitors, either FDA approved or in clinical trials (**Table S3**). This manually curated target-drug(s) database is dominated by signaling proteins and established oncoproteins, including c-KIT. Pharmacological agents with narrow therapeutic indices—such as those targeting neurotransmitters, ion channels, and vasoactive drugs—were purposefully removed from the database as they are less likely to be successfully repurposed in oncology.

*OncoTarget* simply analyzes the [meta]VIPER outputted protein activity measurements for these 180 actionable proteins, and provides a multiple-testing corrected significance value for the corresponding NES. We use a conservative threshold (BH-corrected *p* < 10^−5^) to identify candidate proteins eliciting essentiality when targeted by a pharmacological inhibitor, for *in vivo* validation. For comparison, *OncoTarget* was run using the dGES (mRNA Z-scores) of samples relative to the centroid, instead of protein activity as its input, and is presented in Figure S5.

### OncoTreat Analysis

We have previously described *OncoTreat*^28,33^ as a methodology to systematically elucidate compounds capable of significantly reversing the activity of the 25↑+25↓ MRs comprising the *tumor checkpoint module* (TCM) that regulates the tumor state in general, or more specifically that which regulates metastatic progression or therapeutic resistance.

For the current study, we adapted *OncoTreat* to identify MR-activity reversing compounds to target the imatinib-resistant state. The requisite steps include:

1. <u>Identifying cell lines (typically 1 or 2) or organoid lines jointly representing high-fidelity (i.e., cognate) *in vitro* models</u> for the samples of interest (e.g., imatinib-resistant LRG GISTs), based on MR-activity conservation (BH-corrected *p* < 10^−5^), as assessed by enrichment analysis, see next section on OncoMatch.
2. Generating RNASeq profiles within each selected cell line, from Step 1, at 24h following perturbation with a library of clinically relevant oncology drugs. We thus generated pre-and post-drug perturbation profiles for both GIST-T1 (RRID:CVCL_4976) and GIST430 (RRID:CVCL_7040), using 46 drugs, including 30 FDA approved antineoplastics and 16 late-stage experimental drugs in phase II or III oncology clinical trials (**Table S5**). GIST-T1 cells were grown in RPMI culture media with 10% FBS, while GIST430 cells were grown in IMDM with 10% FBS plus low dose imatinib 100nM to maintain relative imatinib resistance, per published recommendations^90,91^. Initial 10-point dose response curves for each drug in each cell line at 48 hours were performed using the CellTiter-Glo assay, to estimate an EC_20_, the concentration at which cell growth is inhibited by 20% compared to vehicle control. Subsequently, cells were harvested for RNA extraction at 24-hours following perturbation with each drug at two sublethal concentrations—the EC_20_ and one tenth of this concentration. The lower dose was introduced to assess the potential pharmacologic window of each drug. Sublethal drug concentrations effectively reveal drug mechanism of action rather than confounding effects associated with cell stress or death pathways^28,57,58^. We also capped concentrations at each drug’s *C*_Max_, defined as the peak serum concentration for the drug’s maximum tolerated dose (MTD), from published pharmacokinetic studies in humans, when available, thus optimizing the translational potential of predicted drugs. DMSO was selected as a universal *in vitro* solvent (vehicle). Multiplexed, low depth (1M to 2M reads) RNASeq profiles were generated using 96-well plates via the PLATESeq technology, using fully automated microfluidics for increased throughput and reproducibility^60^. Eight DMSO-treated controls were included in each plate, to avoid plate-dependent batch effects and to mitigate the inherent variability of DMSO treatment.
3. <u>Generating a subproteome-wide context-specific Drug Mechanism of Action (MoA) for each drug</u>, as represented by the differential activity of each protein in drug- vs. vehicle control (DMSO)-treated cells. Differential protein activity was assessed by VIPER analysis, see above.
4. <u>Identifying tumor-specific candidate MRs and the MR-activity signature they comprise</u>, by metaVIPER analysis of the sample’s dGES, using the opposing cluster of LRG samples as the *reference model*.
5. Finally, prioritizing pharmacological agents based on the statistical significance of the enrichment of the tumor sample’s MR-activity signature (*i.e.*, 25↑+25↓ MRs) in proteins inactivated and activated in drug- vs. DMSO-treated cells, respectively, with negative NES indicating MR-activity reversal (BH-corrected *p* < 10^−5^, 1-tailed aREA). The number of candidate MR proteins (n=50) used to assess MR-activity reversal, which for this step is restricted to only transcription factors and co-factors, was selected because we have shown that, on average, across all of TCGA, the vast majority of functionally-relevant genomic events can be found upstream of the top 50 VIPER-inferred candidate MR proteins^70^. We also demonstrated in^28^ that a wide range in the number of candidate MRs included in the MR-activity signature has minimal effect on the ordering of *OncoTreat* predictions for drug sensitivity.

We have subsequently completed an expanded screen in GIST430, using 333 drugs, including 122 FDA approved antineoplastics, 192 late-stage experimental drugs in phase II or III trials, and 19 compounds from diversity libraries presenting cell line-specific EC_50_ ≤ 2 μM (**Table S6)**. Data from the expanded screen were not yet available for the original *OncoTreat* predictions for *in vivo* validation, but are shared here, including an unsupervised clustering of drug mechanism based on induced differential protein activity (**Figure S4**). For expanded screens, we have attempted to be all inclusive, with the exception of therapeutic antibodies and antibody drug conjugates due to a lack of availability and immunotherapy agents due the inability to appropriately characterize their intended effects *in vitro*. Additional miscellaneous compounds with EC_50_ ≤ 2μM in the selected cell line were also included. Most compounds were purchased from SelleckChem or Tocris.

### OncoMatch, Cell Line and Patient-Derived Xenograft (PDX) Model Fidelity Analysis

Model fidelity was assessed based on the statistical significance of MR-activity conservation between cluster ***C_1_*** (imatinib-resistant) LRG GIST tumors and a model-derived sample.

For the purposes of OncoMatch analysis to cell line models only, protein activity was computed by generating a dGES comparing each cell line against a large repository of cancer cell lines (*reference model*), which includes both the Cancer Cell Line Encyclopedia (CCLE)^92^ and the Genentech Cell Line Screening Initiative (gCSI)^93^. Meanwhile, protein activity for LRG tumors was computed against an analogous *reference model* — RNA profiles of all tumor samples in TCGA. Lineage matched normal or adjacent tissue is not used as a *reference model* for OncoMatch, as regulators of differential cell proliferation signaling of tumor v. normal dominates the resulting MR signature. Next, we computed the enrichment of the MR-activity signature (25 most active and 25 most inactive patient tumor-specific MRs) in differentially active and inactive proteins in the model (OncoMatch). The aREA^32^ test was again used, but any suitable enrichment analysis algorithm could be substituted. All cell lines used in these analyses were re-sequenced on-site, to capture any potential drift effects. A threshold of BH-corrected *p* < 10^−5^ was used to identify at least moderate-fidelity models. The analysis was used to evaluate the suitability of the GIST430 and GIST-T1 models for the generation of perturbational profiles that optimally capture the effects of tested drugs on the activity of MR proteins.

We worked with Crown Bioscience to characterize 18 of their existing GIST patient-derived xenograft (PDX) mouse models and identified a small subset that was remarkably representative of imatinib-resistant GIST. For the purposes of OncoMatch analysis between patient tumors and PDX models, protein activity was computed by first generating a dGES relative to the centroid of the LRG GIST and Crown Bioscience PDX datasets, respectively, followed by metaVIPER analysis and application of aREA to evaluate MR-activity signature conservation. We thus evaluated the fidelity of PDXs prior to testing drug predictions, and further confirmed that those drug predictions were also concordant for the PDX tumor before proceeding.

### Therapeutic drug testing

All *in vivo* studies were performed by Crown Bioscience. Initial maximum tolerated dose (MTD) studies were performed at 3 dose levels for imatinib, regorafenib, and the six candidate drugs in non-tumor bearing NOD/SCID mice. Reported doses in the literature for PDX studies were used as a starting point for the dose levels evaluated. Dosing continued for 14 days, with an additional 7 days of post-treatment monitoring for toxicity. Clinical observations were performed twice weekly and body weight measurements daily or every other day at minimum.

For the efficacy studies in *GS5106* and *GS5108*, tumor fragments from stock mice were harvested and used for inoculation into mice. Mice were ear tagged. Each mouse was inoculated subcutaneously in the right front flank with a specific PDX tumor fragment (3×3×3 mm). Implanted mice were monitored for palpable tumors, or any changes in appearance or behavior. Daily monitoring took place for mice showing any signs of morbidity. Once tumors were palpable, tumors were measured twice weekly using calipers. Tumor volume was estimated using the following equation: (longest diameter * shortest diameter^2^)/2.

Once average tumor volume reached 150-250mm^3^, mice were randomly assigned to the respective treatment groups and dosed on day 1. Dosing continued for up to 21 days or until death/sacrifice, with an additional 7 days of observation post-treatment. Tumor volume was measured twice weekly for the duration of the study. One individual was responsible for tumor measurements for the duration of the study. Body weight was also measured at least twice weekly for the duration of the study, and frequency of monitoring was increased to every other day when toxicity was noted. If body weight loss of >10% was observed, Dietgel was given *ad libitum*, and body weight loss of >15% triggered a dosing holiday until weight loss recovered to <5%. If body weight loss of >20% was observed, the animal was monitored daily for signs of recovery for up to 72 hours. If there were no signs of recovery, the animal was sacrificed as per Crown Bioscience’s IACUC protocol regulations.

The DRAP package for R^62^ was used to plot tumor volume curves and compute mean tumor growth inhibition (TGI) on treatment (21-days) for each drug arm versus vehicle control. TGI was calculated using the formula 1 – F(Vt)/F(Vc), where F(Vt) is the difference in mean tumor volume from baseline in the treatment arm, and F(Vc) is the difference in mean tumor volume from baseline in the vehicle control arm at each timepoint. TGI ≥ 1.0 (i.e., ≥ 100%) indicates that mean tumor volume has regressed from baseline in the treatment arm, while a negative value, TGI < 0 (i.e., a negative percentage), indicates that the tumor volume increase in the treatment arm exceeded that seen in the control arm.

The linear mixed model for repeated measures^62,94^, as implemented in DRAP, was used for statistical significance testing to determine if a drug significantly impacted tumor growth from time 0 to the EoT timepoint. For both mean TGI and statistical significance testing, drugs were compared to the concurrent vehicle control arm unique to the five separate parts of the PDX therapeutic study as presented in **Figure 5**.

### Pharmacodynamic (PD) Assessments of MR-activity reversal

Samples for PD assessment were procured from two mice per treatment arm in *GS5106* and *GS5108*. Mice were randomly selected for early sacrifice, independent of tumor size, three hours following the third dose, and were excluded from response assessment. We performed RNASeq and subsequent VIPER on paired drug vs. vehicle control-treated PDX tumor samples. MR-activity reversal was assessed based on the statistical significance of the enrichment of the MR-activity signature (i.e., 25↑+25↓ MRs of the baseline PDX tumor profile) in proteins inactivated and activated in drug vs. vehicle control-treated mice, respectively, again using aREA, although alternative enrichment tests may be used. Negative NES indicates MR-activity reversal (BH-corrected *p* < 10-5), analogous to *OncoTreat*. We also assessed PD reversal of the integrated ***C_1_*** imatinib-resistant cluster MR-activity signature, as presented in **Figure S6**.

### Data Availability

RNASeq raw data of 34 LRG patient tumors and RNASeq profiles of on-treatment (pharmacodynamic) PDX tumor samples generated as part of this work have been deposited to GEO. RNASeq data of 18 established GIST PDX models were obtained via a data use agreement with Crown Bioscience. Relevant high-throughput drug perturbation data used for *de novo* assessment of drug MoA are available on Figshare (FS: https://figshare.com/articles/dataset/Drug_Perturbation_Profiles_in_GIST_Cell_Lines/30247381). The version of TCGA and TARGET processed data (RNASeq counts) used to generate networks is available on FS (https://figshare.com/articles/dataset/TCGA_and_TARGET_RNASeq_Counts_Data_EntrezID_/30247285). The ARACNe-generated networks used to run VIPER are also available on FS (https://figshare.com/articles/dataset/Available_Interactomes_Networks_/30246550). Additional requests should be directed to the corresponding author.

## Acknowledgements

This work was supported by the NCI Office of Cancer Target Discovery and Development (CTD2) award U01CA272610, the NCI Outstanding Investigator award R35CA197745, and the NIH Shared Instrumentation Grant S10OD032433, all to A. Califano. Also, this publication was supported in part through the National Cancer Institute Cancer Center Support Grant P30CA013696 to CUIMC and by the National Center for Advancing Translational Sciences, National Institutes of Health, through Grant Number UL1TR001873. The content is solely the responsibility of the authors and does not necessarily represent the official views of the NIH.

## Supplementary Figures and Tables

**Figure S1.** Pathway enrichment analysis on the integrated protein activity signature of all cluster ***C*_1_** (imatinib-resistant) vs. ***C*_2_** tumors. The MSigDB gene set collections for Gene Ontology, Cancer Hallmarks, Immunological Signatures, and Oncogenic Signatures are used for this analysis, with enrichment for all genes in a pathway assessed in the full ranked protein activity signature of each cluster, as inferred by metaVIPER. As the cluster specific signatures were computed iteratively, pathways enriched in ***C*_1_** tumors (dark red) are [inversely] de-enriched in ***C*_2_** tumors, and pathways enriched in ***C*_2_** tumors (cyan) are de-enriched in ***C*_1_**tumors. The analysis suggests imatinib resistance is associated with a complex cellular reprogramming affecting transcriptional machinery, DNA repair, oxidative phosphorylation, immunological signaling, and MYC proliferative signaling, amongst others.

**Figure S2.** UMAP projection of metaVIPER inferred protein activity signatures of scRNA profiles of high-risk GIST, annotated by c-KIT activity and KIT mRNA differential expression. (***Top A-I***) The same UMAP projections of metaVIPER inferred protein activity signatures (n=7,070 proteins) of malignant compartment single cells displayed in **Figure 2** are re-created here, but with individual cells now annotated by differential activity of c-KIT. Larger red dots represent cells with higher c-KIT activity, expressed in NES, while larger blue dots represent cells with decreased c-KIT activity (negative NES). As in **Figure 2**, the two tumors without clinical progression on imatinib and without any treatment-emergent mutations associated with resistance are in panels ***A,B***; the four tumors without established clinical progression but with mutations associated with the development of resistance are in panels ***C,D,E,F***; and the three tumors with confirmed clinical progression on imatinib are in panels ***G,H,I***. (***Bottom A-I***) Same as top, except individual cells are now annotated by differential *KIT* mRNA activity, expressed as Z-score.

**Figure S3.** Inference of copy number variation from scRNA profiles of high-risk GIST, annotated by enrichment for the imatinib-resistant cluster signature (ImResS) in putative malignant cells. InferCNV^88^ was used to identify inferred copy number variations, using cells annotated as immune system cells by SingleR^85^ (T cells, NK cells, Monocytes, and Macrophages) as reference. Cells that express the highly specific GIST marker^87^ DOG-1 (also known as ANO1) and also annotated by SingleR as either Smooth muscle cells, Tissue stem cells, Fibroblasts, Chondrocytes, Mesenchymal stem cells, Neurons, or Neuroepithelial cells are entered as potential malignant cells. (A-E) Panels A,B provide InferCNV analysis for two tumors without established clinical progression but with mutations associated with the development of resistance and with sizable ImResS+ subpopulations (***Figure 2C,E***), while panels C-E provide InferCNV analysis for the three tumors with confirmed clinical progression on imatinib (***Figure 2G-I***). Cells that were ultimately characterized as the malignant compartment within each sample based on the above criteria and the InferCNV analysis, are further annotated by their ImResS enrichment, using a small annotation bar in the lower left corner of each panel.

**Figure S4.** Heatmap for the 24-hour expanded drug perturbation screen in GIST430 and the more limited screen completed in GIST-T1. The heatmap shows the differential protein activity profile of 333 drugs in GIST430 cells and 46 drugs in GIST-T1 (rows) compared to vehicle control. Drug profiles are annotated by the cell line in which they were generated and by the drug’s canonical mechanism and the sublethal EC_20_ concentration at which each drug was screened. The 524 VIPER monitored proteins that demonstrate the highest variance in activity across perturbations are shown in the columns, and we use unsupervised hierarchical clustering to organize groups of drugs that induce similar transcriptional response, i.e., context-specific mechanism of action.

**Figure S5.** Consensus clustering and drug predictions for Life Raft patient GIST tumor samples, using differential gene expression analysis alone. (**A**) Unsupervised consensus clustering of tumors was performed on the differential gene expression signatures (dGES) of each sample—computed as Z-scores relative to the full dataset’s centroid—using a partition around medoids (PAM) approach, with n=10,000 iterations. The cluster solution for k=5 partitions is shown. (**B**) Heatmap representing the most differentially overexpressed of a panel of 180 directly druggable proteins based on mRNA Z-score. Columns are organized in the same order as ***Figure 1A*** and color intensity corresponds to the −log10 (BH-corrected *p*) computed from the mRNA Z-score. The prior cluster assignment (***Figure 1***) and clinical annotation are provided on top of the heatmap. **(C)** Drug sensitivity predictions based on reversal of the most differentially expressed genes at the mRNA level for each tumor sample, in the drug signature generated in the GIST430 model. Enrichment was assessed by the aREA algorithm, with negative normalized enrichment scores (NES) indicating reversal of the differential gene expression signature, and the associated lower-tail *p*-value. Color intensity in the heatmap corresponds to the −log10 (BH-corrected *p*), and the predictions are clustered by tumor (*columns*) and drug (*rows*), with cluster reliability indices for cluster assignment shown as barplots on the top and right-hand side of the heatmap. The prior cluster assignment and clinical annotation are provided at the bottom of the heatmap.

**Figure S6.** Pharmacodynamic assessment of PDX MR-activity reversal. (**A**) Pharmacodynamic change induced by each treatment arm in the activity of the top 25 activated and top 25 inactivated MRs, as defined from the baseline PDX profiles, which corresponds to the red and blue hit marks on the enrichment plots in ***Figure 6A-C***. (**B**) Pharmacodynamic change in activity of the top 25 activated and top 25 inactivated MRs of the integrated signature of all cluster C_1_ (imatinib-resistant) patient tumors. Reversal of this signature based on *in vitro* assessment formed the initial basis for all OncoTreat predictions, and *in vivo* recapitulation of this effect may be a requisite for drug efficacy for *OncoTreat*-predicted drugs.

**Table S1.** Quality control metrics on Life Raft patient GIST samples, including tumor cell percentage on hematoxylin and eosin (H&E) slide review by a clinical pathologist, RNA concentration, library prep used, and mapped reads and detected genes from bulk RNASeq.

**Table S2.** Extended demographic and clinicopathologic annotation for all Life Raft samples used in the downstream analyses, including *KIT/PDGFRA* genotype, relative timing of sample acquisition, time on various treatments, and progression and survival outcomes when available. Samples from top to bottom are presented in the same order as they appear in the clustering heatmap (***Figure 1A*** from left to right).

**Table S3.** Curated list of proteins with high-affinity inhibitor drugs for OncoTarget analysis.

**Table S4.** Demographic and clinicopathologic annotation for all 9 samples from which scRNA profiles were generated, as provided in gene expression omnibus entry GSE54762 and Liu et al.^44^.

**Table S5.** List of drugs in the initial GIST430 and GIST-T1 perturbation screens. The EC_20_ and lower concentration at which each drug was tested is reported, along with FDA approval status.

**Table S6.** List of drugs in a subsequent expanded GIST430 perturbation screen. The EC_20_ at which each drug was tested is reported, along with FDA approval status.

**Table S7.** Animal-level data across study duration for tumor volume, medication administration record, and body weight, with deaths highlighted in red. Each sheet corresponds to the 5 study parts shown in ***Figure 5A-E***.

**Table S8.** Detailed summary of therapeutic response at end of treatment time point, organized by PDX model and separate parts of the study performed with independent control arms. The presented t-statistic and *p*-value are from a linear mixed model accounting for repeat measures over time of tumor volume, as implemented in the DRAP package.

