## Supplementary figures and images for "Systematic elucidation and pharmacologic targeting of non-oncogene dependencies in imatinib-resistant gastrointestinal stromal tumor"

### Figure S1

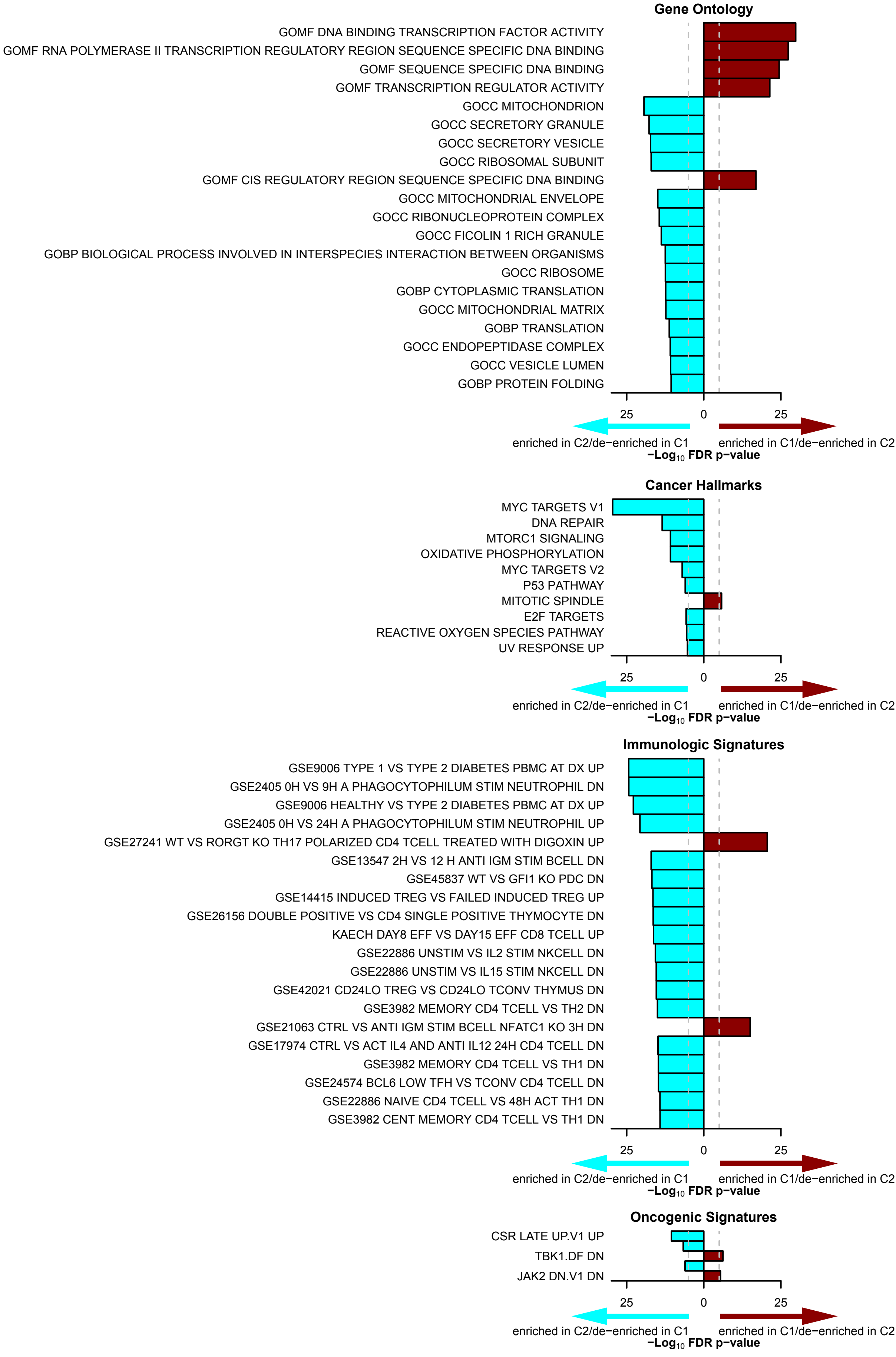

### Figure S2

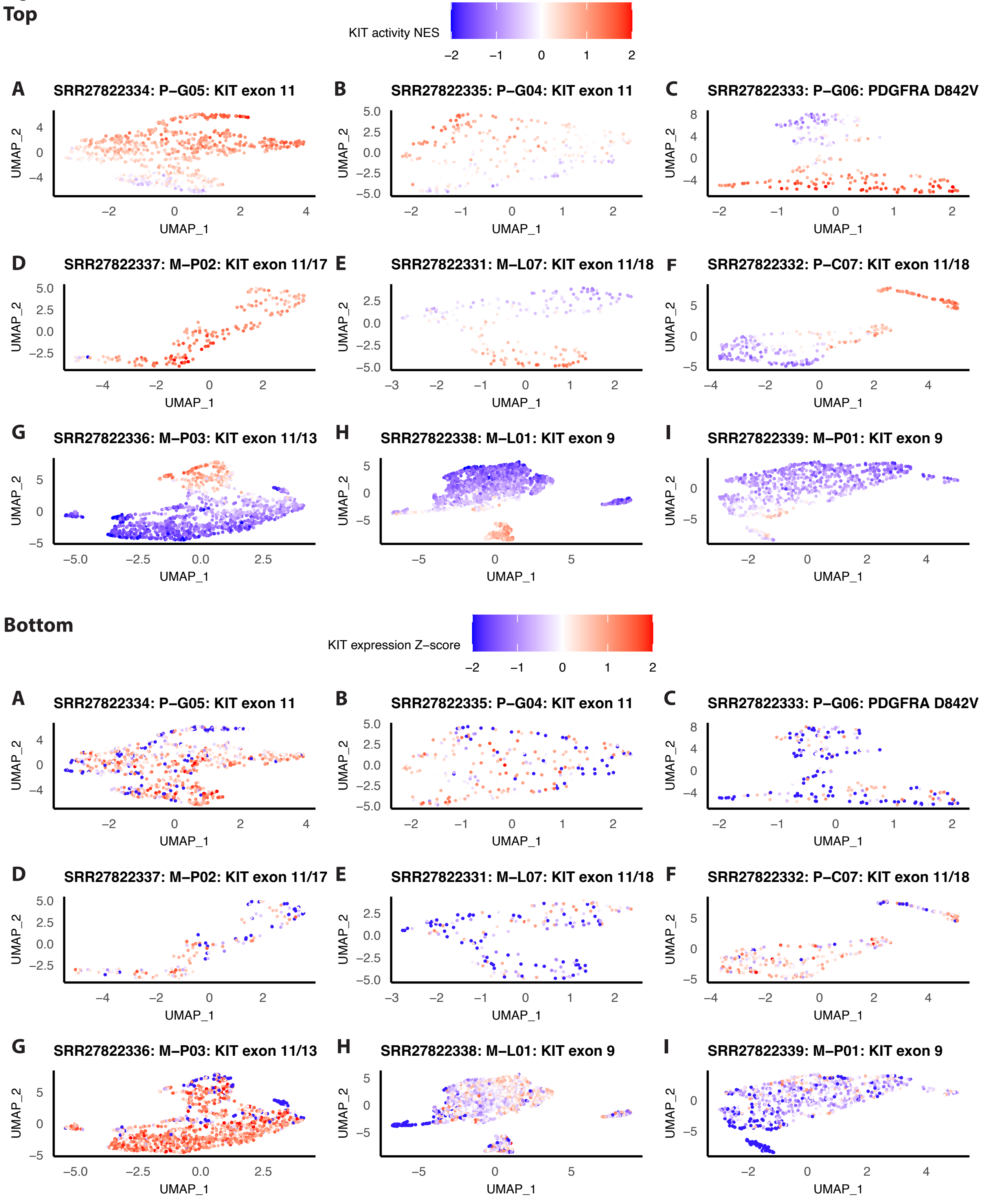

### Figure S3

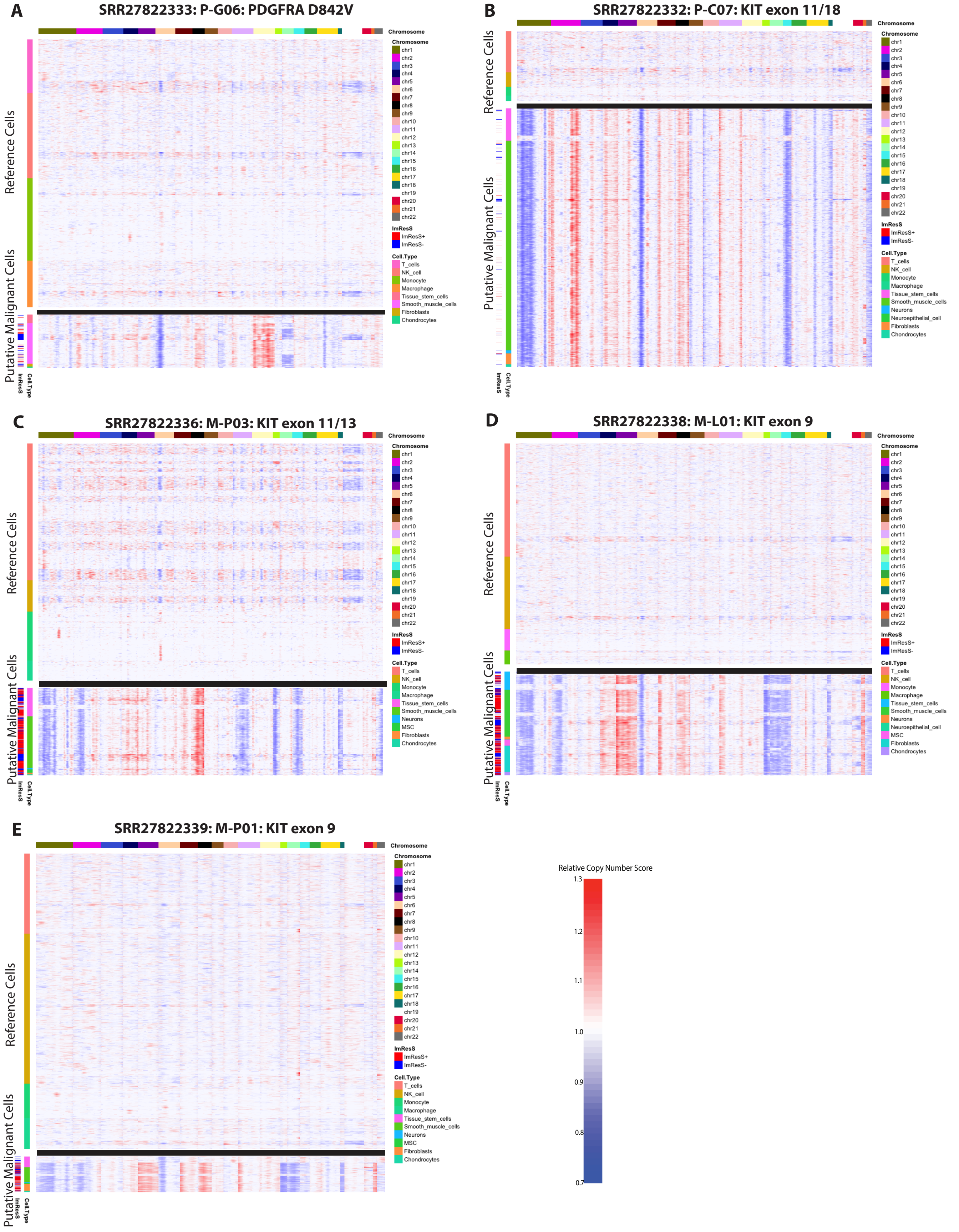

### Figure S4

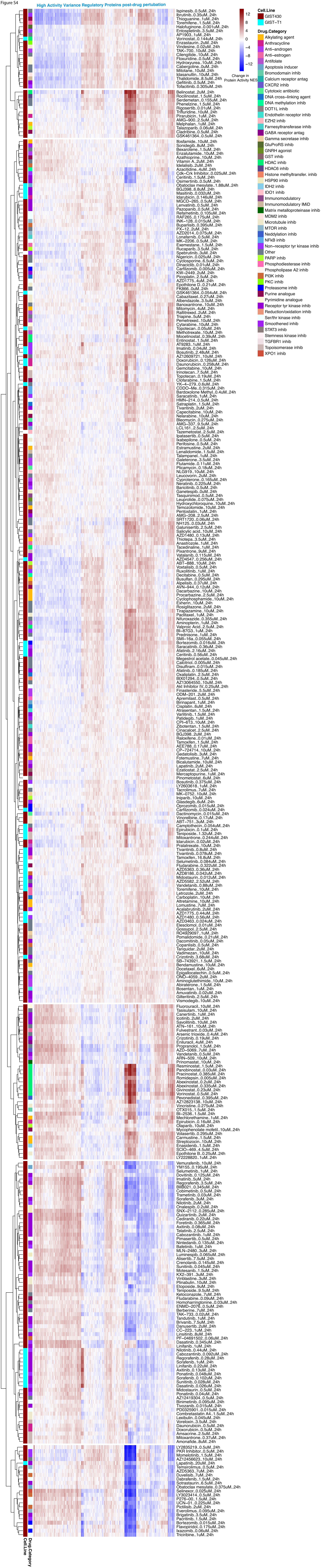

### Figure S5

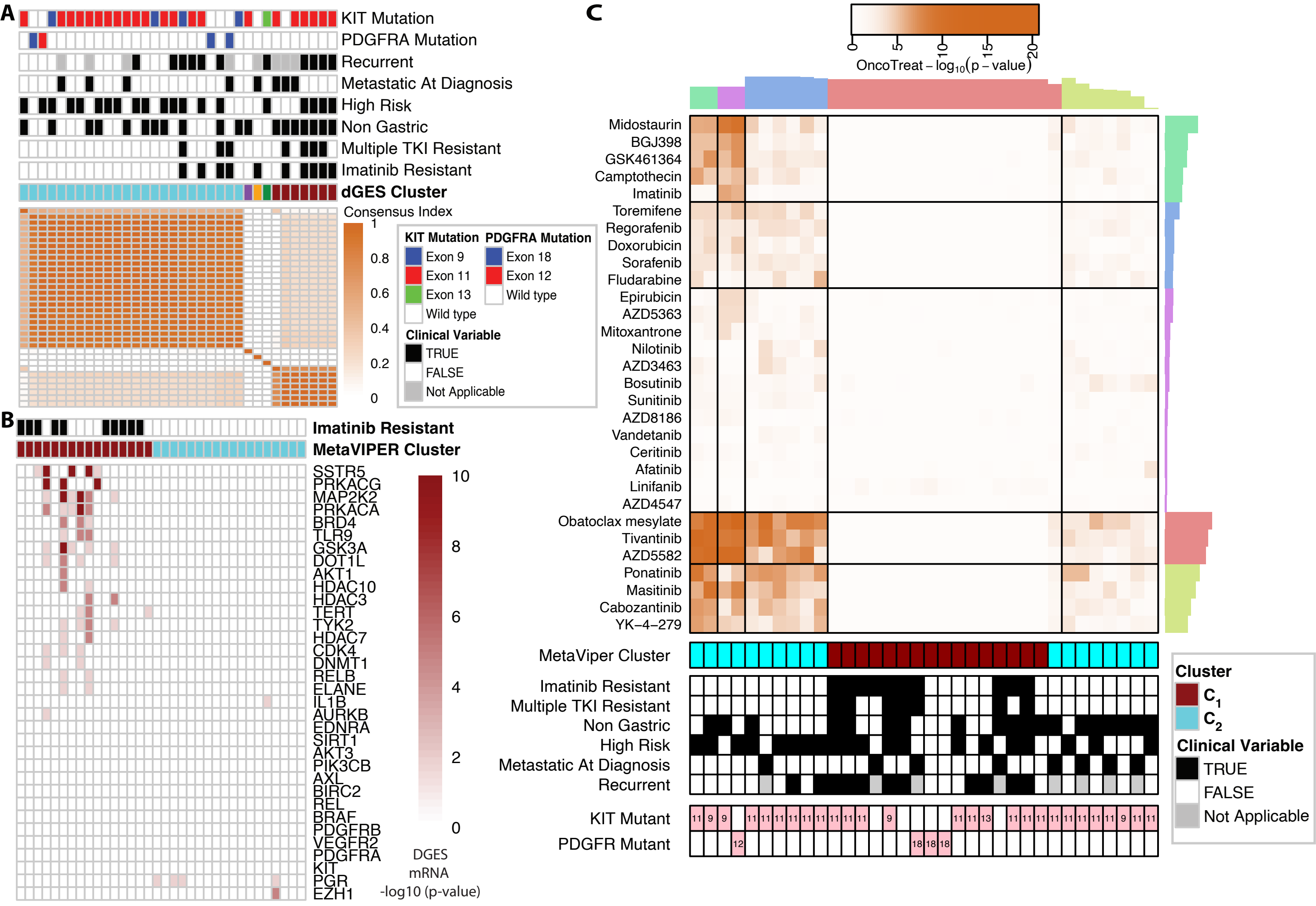

### Figure S6

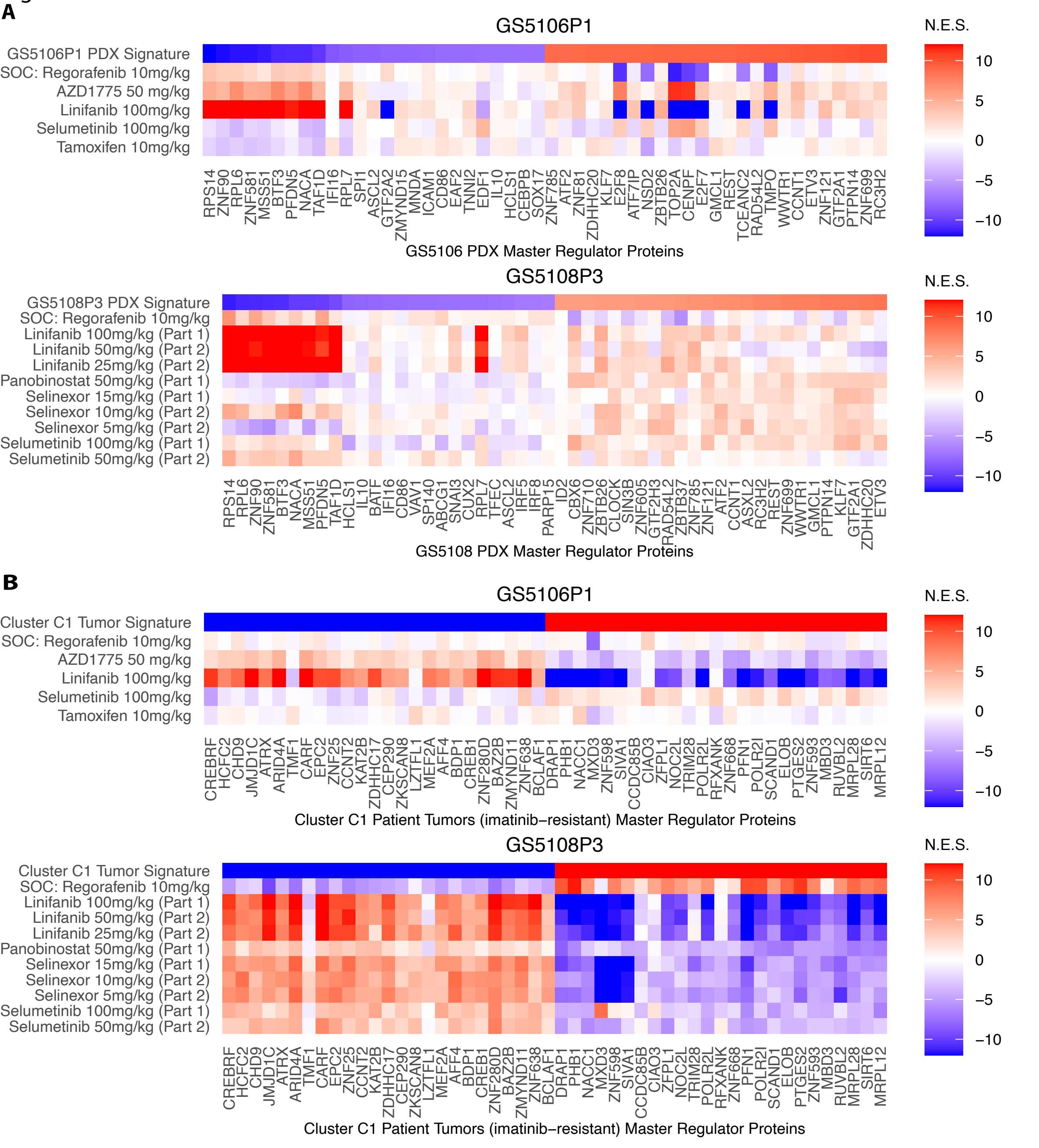
